# Structural basis and physiological significance of non-canonical Gs coupling to the prototypical Gi-coupled melatonin MT_1_ receptor

**DOI:** 10.1101/2025.11.08.687348

**Authors:** Atsuro Oishi, Hiroyuki H. Okamoto, Keisuke Ikegami, Ronan McHugh, Bernard Masri, Tsukasa Kusakizako, Kazuhiro Kobayashi, Angeliki Karamitri, Erika Cecon, Julie Dam, Miki Nagase, Irina G. Tikhonova, Osamu Nureki, Ralf Jockers

## Abstract

G protein-coupled receptors (GPCRs) transduce extracellular stimuli into intracellular signals by coupling to various heterotrimeric G proteins. However, the rules governing G protein preference remain largely elusive. MT_1_ and MT_2_ are prototypical G_i/o_-coupled GPCRs responding to melatonin, a hormone secreted in a circadian manner. We show here that MT_1_, but not MT_2_, couples also to G_s_ proteins *in vitro* and activates the G_s_/cAMP pathway upon long-term melatonin exposure *in vivo*, mimicking physiological dawn conditions. We solved the cryo–electron microscopy structure of the melatonin-MT_1_-G_s_ complex at 3.0Å resolution, which revealed a strikingly distinct binding mode compared to the MT_1_–G_i_ complex. The third intracellular loop of MT_1_ emerges as a key stabilizer for G_s_ coupling, a feature previously unrecognized. This is the first solved receptor-G_s_ complex of a primary G_i_-coupled GPCRs, providing new structural and functional insights into G protein selectivity and circadian switch of G protein coupling.

## Introduction

G protein-coupled receptors (GPCRs) represent the largest family of membrane proteins involved in cellular signal transduction. GPCRs activate their primary transducers, heterotrimeric G proteins, that are classified in four families: G_s_, G_i/o_, G_q/11_, and G_12/13_. Among those, G_s_ and G_i/o_ proteins have opposing roles as they are stimulating or inhibiting adenylyl cyclases (AC) and cyclic AMP production, respectively. Large-scale common G protein coupling datasets show that a particular GPCR can couple to one or several G protein types^1^. G_s_-coupled GPCRs show the lowest coupling promiscuity and rarely co-couple to G_i/o_ proteins or with a large difference in their strength of activation^1^. The few exceptions of G_s_/G_i/o_ co-coupling include the primarily G_s_-coupled 5-HT4, EP4 and ß1AR^2–4^, as well as the primarily G_i/o_-coupled GPR139^5^.

MT_1_ and MT_2_ are high-affinity receptors for melatonin (5-methoxy-N-acetyltryptamine) that primarily couple to G_i/o_ proteins, leading to inhibition of cAMP production and the PKA/CREB pathway, activation of ERK1/2, and regulation of ion channels^6–8^. The molecular structures of MT_1_ and MT_2_ receptors have been solved, all in the presence of synthetic agonists, with some in complex with the G_i_ protein^8–13^. The stimulation profile of melatonin receptors is unique, governed by the circadian secretion pattern of melatonin in the pineal gland with peak levels during the night^14^. Due to this rhythmic pattern of melatonin secretion, melatonin receptors regulate and synchronize a variety of physiological functions including sleep/awake rhythm, seasonal reproduction, and retina physiology, with currently marketed drugs indicated for sleep and circadian disorders and major depression^15,16^. Whereas most effects of melatonin receptors appear to be transmitted by G_i/o_ proteins, coupling to G_s_-dependent pathways has been suggested for MT_1_^17–19^. However, whether this is due to direct G_s_ coupling or to downstream crosstalk is unknown. The physiological relevance of dual G_i_/G_s_ coupling, with apparently opposing signaling outcomes, also remains elusive and intriguing. Here, we provide functional evidence for G_s_ coupling to MT_1_ under physiologically relevant conditions in the mouse hypophysial *pars tuberalis* (PT), and report the cryo-electron microscopy (cryo-EM) structure of the MT_1_–G_s_ complex in the presence of the natural ligand melatonin. The unique interaction mode of G_s_ depends on the third intracellular loop of MT_1_ (ICL3) which is required to stabilize the interaction with MT_1_.

## Results

### Melatonin MT_1_ receptor activates both the inhibitory G_i_/cAMP and the stimulatory G_s_/cAMP pathway

Acute stimulation with melatonin of HEK293 cells expressing low levels of human MT_1_ receptor inhibited forskolin (FSK)-stimulated cAMP production with an pEC_50_ of 9.96±0.34 (n=6), as expected for this prototypical G_i/o_-coupled receptor (Fig. 1a). In contrast, cells expressing high MT_1_ levels showed a biphasic behavior with inhibition of cAMP production at low melatonin concentrations (pEC_50_=10.3±0.35; n=6) and stimulation of cAMP production at higher melatonin concentrations (pEC_50_=8.6 ±0.66; n=6) (Fig. 1a). Similar experiments in cells expressing high levels of the closely related G_i/o_-coupled human MT_2_ melatonin receptor showed only the inhibitory component (pEC_50_=9.93±0.23; n=6), indicating that the biphasic behavior is a unique feature of MT_1_.

**Fig 1.**
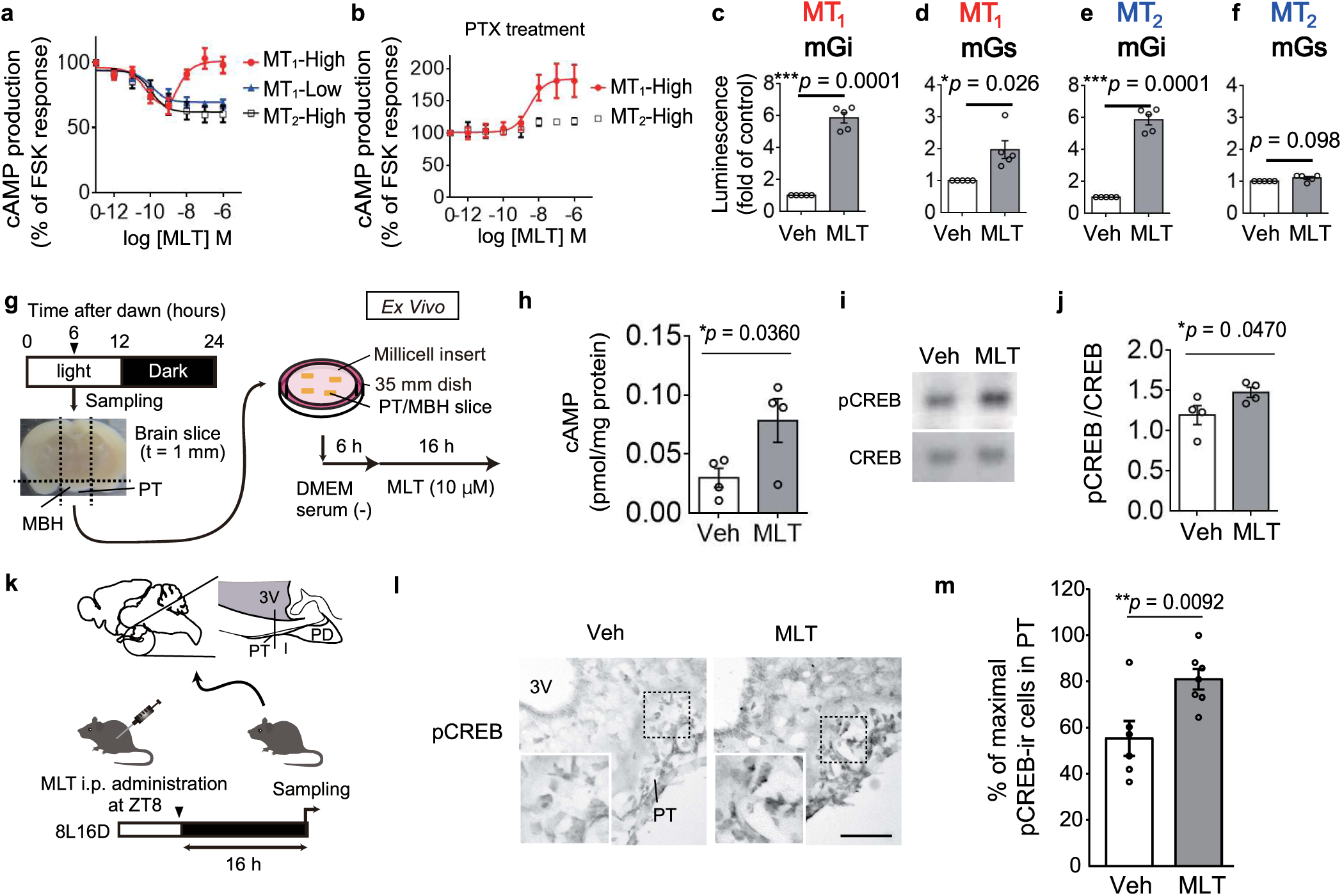
Melatonin MT_1_ receptor activates the stimulatory G_s_/cAMP pathway *in vitro*, *ex vivo* and *in vivo*. (a-f) Melatonin MT_1_ receptor activated G_s_-cAMP pathway in HEK293T cells. (a) (b) Concentration–response effect of melatonin on forskolin (FSK)-induced cAMP production (30 min, FSK [2 µM], IBMX [1 mM]) in HEK293T cells expressing MT_1_-WT or MT_2_-WT, assessed by HTRF assay without (a) or with (b) PTX treatment (400 ng/mL, 4 h). Data represent means ± SEM of at least 4 independent experiments. (c-f) NanoBiT complementation assay showing recruitment of LgBiT–miniG_i_ (c,e) or LgBiT–miniG_s_ (d,f) to human MT_1_-NP (c, d) or MT_2_-NP (e, f) upon melatonin stimulation (1 µM, 2 min). Data are presented as means ± SEM of five independent experiments. Statistical significance was determined using paired two-tailed t-tests, exact p-values are reported. (g-j) Melatonin (MLT) induces cAMP production in mouse pituitary *pars tuberalis* (PT) *ex vivo*. (g) MLT (10 µM) were exposed to sliced PT / mediobasal hypothalamus (MBH) tissue complex of B6 mice kept under 12L/12D conditions. (h) Sixteen-hour exposure of MLT significantly increased cAMP levels in the PT/MBH compared with control 0.1 % DMSO exposure group. Data are means ± SEM of 4 independent experiments. (i) Western blot analysis of phosphorylate CREB (pCREB [Ser133]) in cultured PT/MBH tissues. MLT significantly increased pCREB. Data are means ± SEM of 4 independent experiments. Unpaired t-tests with Welch’s correction were used and one-tailed p-values are reported. (k-m) MLT induces cAMP production in mouse pituitary PT *in vivo*. (k) MLT (0.26 mM / 0.1 mL) was injected intraperitoneally (i.p.) at ZT8 to B6 mice kept under 8L/16D conditions. After 16 h, brains were extracted and sectioned coronally for immunohistochemistry. ZT, Zeitgeber time; ZT1 indicates light onset time. (l) Representative images of pCREB immunoreactivity (ir) in the PT and MBH of MLT-injected mice. 3V, Third ventricle. (m) Relative changes of pCREB-ir cells normalized by area in the PT. MLT injection increased pCREB (\*\**p* < 0.01, t-test). Data are means ± SEM of 6-7 independent experiments. Unpaired t-tests with Welch’s correction were used and one-tailed p-values are reported.

Pretreatment of HEK293 cells expressing high levels of MT_1_ and MT_2_ with pertussis toxin (PTX), a G_i/o_ inhibitor, abolished the inhibitory effect of melatonin for both receptors, while the stimulatory effect was preserved for MT_1_ with a pEC_50_ of 8.51±0.38; n=6) (Fig. 1b), which was similar to the value observed in the absence of PTX. This indicates that MT_1_ and MT_2_ couple to PTX-sensitive G_i/o_ proteins and MT_1_ to an additional PTX-insensitive G protein, possibly the stimulatory G_s_ protein. To test this hypothesis, we measured G_i_ and G_s_ coupling directly to MT_1_ and MT_2_ with the nanoluciferase (Nluc) complementation assay^20,21^. MT_1_ and MT_2_ were fused at their C-terminus to the NP fragment of Nluc (MT_1(2)_-NP) and coexpressed with miniGα_s_ or miniGα_i_ proteins fused to the LgBiT fragment of Nluc (LgBit-miniGα_s/i_) to monitor G protein recruitment through Nluc complementation^16,17^. Melatonin-promoted recruitment was observed between LgBit-miniGα^i^ and both receptors (Fig. 1c,e) and between LgBit-miniGα^s^ and MT^1^-NP, but not MT^2^-NP (Fig. 1d,f), indicating that MT^1^ couples to G^i^ and G^s^ while MT^2^ couples only to G^i^. Consistently, bi-vs monophasic melatonin concentration-response curves of FSK-stimulated cAMP production were also recapitulated with NP-tagged human MT^1^ and MT^2^, respectively (Supplementary Fig. S1a,b). Similar results were obtained for mouse MT^1^ and MT^2^ indicating that mice could be a relevant model for in vivo validation (Supplementary Fig. S1c,d).

MT^1^ is abundantly expressed in the rodent pituitary PT whereas MT^2^ is absen^t22,23^. Acute melatonin treatment has been shown to inhibit FSK-stimulated cAMP production in cultured PT cells^24^. To simulate physiologically relevant long-term stimulation conditions, we first established ex-vivo cultures from mouse PT and stimulated them for 16h with melatonin (Fig. 1g). Under these conditions, melatonin promoted cAMP production (Fig. 1h) and further downstream CREB phosphorylation (Fig. 1i,j). Intraperitoneal administration of melatonin increased CREB phosphorylation in the mouse PT 16h after treatment (Fig. 1k-m). Collectively, these data indicate that endogenously expressed MT_1_ couples to the G_s_/cAMP pathway in vivo when stimulated for 16h mimicking the end of the dark phase under short-day conditions.

### Structural insights into the MT1-miniGs signaling complex

To obtain structural evidence for G_s_ coupling to MT_1_, we determined the structure of the MT_1_-miniG_s_ complex by cryo-EM at 3.0 Å global resolution, together with the endogenous ligand melatonin, Gβ_1_, Gγ_2_ subunits and the Nb35 fragment (Fig. 2a, b). The overall structure of MT_1_ is similar to the MT_1_ structure in the previously reported MT_1_-Gi complex^12^. In the MT_1_-miniG_s_ complex, MT_1_ is in an active state, supported by the structural similarity at the conserved activation motifs within the MT_1_-G_i_ complex (Extended Data Fig. 3a-e). At the G_s_ interface, the MT_1_-miniG_s_ displayed a novel binding mode, where Arg125^3.50^, in the DRY motif of MT_1_ made no interaction with Tyr391^G.H.5.23^ of G, while this interaction is observed in almost all of the reported structures of class A GPCR-G_s_ complex (Fig. 2c). Instead of the typical interaction between Arg^3.50^ and Tyr391^G.H.5.23^, Tyr128^3.53^ at TM3 of MT_1_ formed a hydrogen bond with Gln390^G.H.5.22^ of G_s_, and Ser132^34.50^ at intracellular loop (ICL)2 of MT_1_ formed a hydrogen bond with His387^G.H.5.19^ of G_s_. In parallel, Leu133^34.51^ at ICL2 of MT_1_ formed a hydrophobic interaction with the pocket of miniG_s_ formed by H41^G.S1.02^, F219^G.S3.03^, F376^G.H5.08^, R380^G.H5.12^ and I383^G.H5.15^, which is observed in most of the G_s_ complexes (Fig. 2c).

**Fig. 2.**
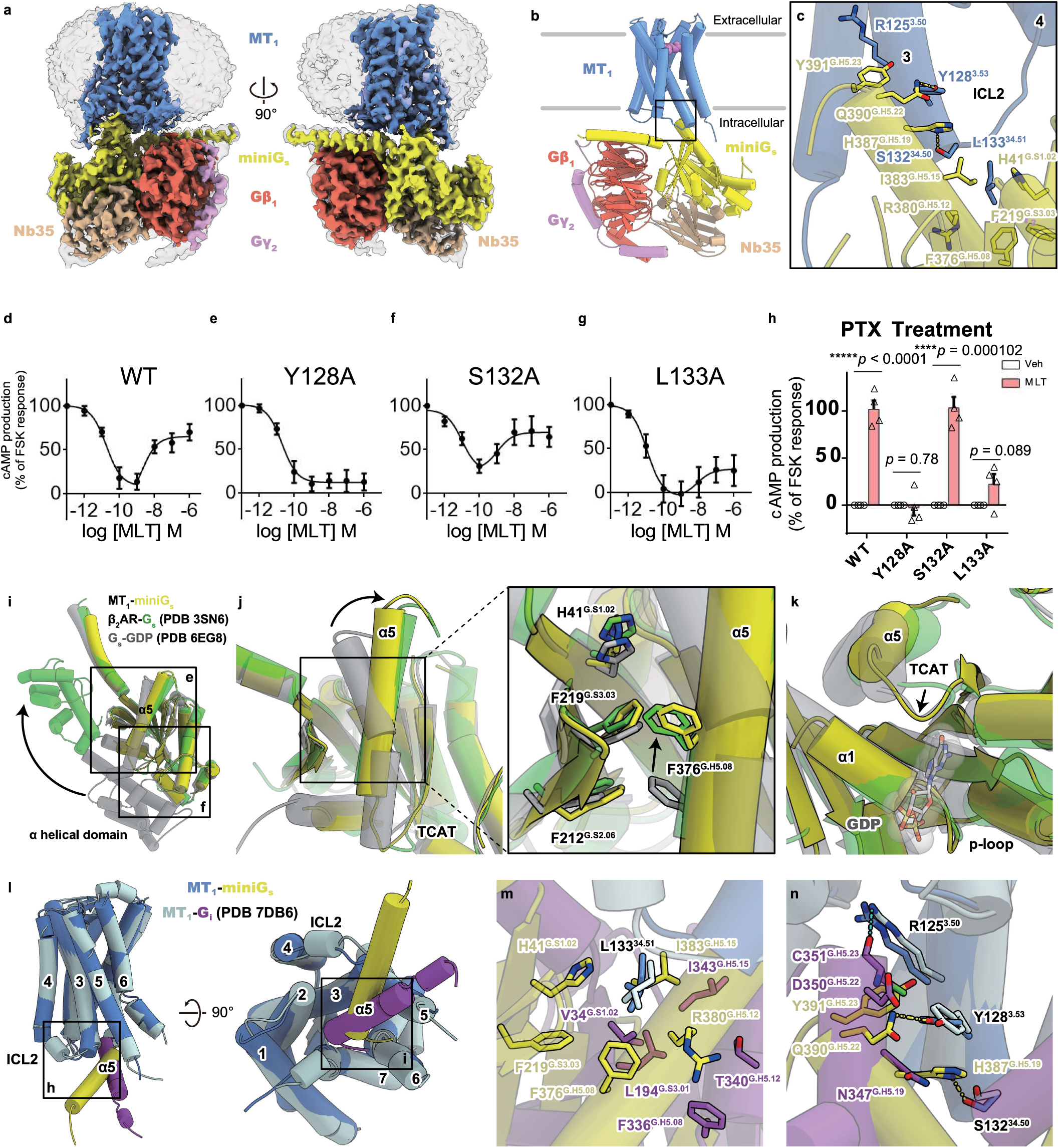
Cryo-EM structure of the MT_1_-miniGs complex. (a) CryoEM density map of MT_1_-miniG_s_ with Gβ_1_, Gγ_2_ and Nb35. MT_1_, miniG_s_, Gβ_1_, Gγ_2_ and Nb35 were shown as blue, yellow, red, pink and brown, respectively. Map including micelle density was shown in transparent style. (b) Overall structure of MT_1_-miniG_s_ with Gβ_1_, Gγ_2_ and Nb35. Melatonin molecule is shown as light purple CPK model. (c) The interface between MT_1_ and miniG_s_. (d–g) Concentration–response effect of melatonin on FSK-induced cAMP production (5 min, FSK (5µM) in HEK293T cells expressing MT1-WT (d) or mutant receptors Y128A (e), S132A (f), and L133A (g), monitored with the BRET-based CAMYEL biosensor. Data represent means ± SEM from six independent experiments. (h) Melatonin (1 µM, 10 min)–induced cAMP production in the absence of FSK in HEK293T cells pretreated with PTX (400 ng/mL, 4 h) and expressing MT1-WT or mutant receptors (Y128A, S132A, L133A) assessed with the BRET-based CAMYEL biosensor. Data represent means ± SEM from four independent experiments. Multiple t-tests with Holm–Šidák correction were used for statistical comparisons, exact p-values are indicated. (i-k) Structural comparison of miniG_s_ protein among MT_1_-miniG_s_ (yellow), β_2_AR-G_s_ (green; PDB 3SN6) and GDP bound G_s_ (gray; PDB 6EG8), focusing on (i) overall structure, (j) α5 helix and (k) TCAT motif. (l-n) Structural comparison with MT_1_-miniG_s_ (light blue and purple; PDB 7DB6), displaying (l) overall structure, (m) ICL2 and (n) TM3.

The importance of these interacting residues was tested by the real-time cAMP assay with the Y128^3.53^A, S132^34.50^A, L133^34.51^A MT_1_ mutants (Fig. 2d-h). All mutated receptor variants showed similar expression levels (Supplementary Fig. S2).. Whereas the inhibitory G_i_-dependent phase was maintained for all mutants, the stimulatory G_s_-dependent phase was completely lost for Y128^3.53^A, not affected for S132^34.50^A and substantially reduced for L133^34.51^A. This suggests the importance of Tyr128^3.53^ and to a lesser extend Ser132^34.50^ in cAMP production via MT_1_-G_s_ coupling, while maintaining MT_1_-G_i_ coupling. Despite this unique binding modes at the MT_1_-miniG_s_ interface, miniG_s_ in the MT_1_-miniG_s_ showed a similar conformation as the previously reported β_2_AR-G_s_ complex (PDB 3SN6) and was distinct from the GDP-bound inactive G_s_ protein structure (PDB 6EG8) (Fig. 2i-k). Phe376^G.H5.08^ moved upward to the C-terminal of the α5 helix in a typical way, causing the extension of α5 helix together with the movement of the TCAT motif at the nucleotide binding site (Fig. 2i-k). From these observations, we concluded that the MT_1_-miniG_s_ complex is a canonical signaling complex rather than an artifact.

Structural comparison with the previous structure of MT_1_ in complex with its primary signal transducer, the G_i_ trimer (PDB 7DB6), revealed that MT_1_ exhibits a distinct binding mode when bound to miniG_s_ (Fig. 2l). While the overall structure of MT_1_ remains similar between complexes, the G protein α5 helixes adopt dramatically different entry angles: G_s_ α5 inserts from the ICL2 side, whereas G_i_ α5 penetrates from the ICL3 side (Fig. 2l). This differential positioning enables unique interactions in each complex. In the MT_1_-G_s_ structure, Leu133^34.51^ forms hydrophobic contacts with the Gα pocket - an interaction absent in the MT_1_-G_i_ complex (Fig. 2m,n). This observation aligns with preserved cAMP inhibition in L133A and Y128A mutants. Conversely, G_i_ residues at equivalent positions do not form similar interactions between Cys351^G.H.5.23^ of G_i_ and Arg125^3.50^ of MT_1_ (Fig. 2n). Critically, G_s_ α5 adopts shallow binding with its C-terminal tip oriented toward ICL2/TM3, while G_i_ α5 penetrates more deeply toward TM5/TM6 regions^12^. We conclude from these observations that the binding mode of the MT_1_-G_s_ complex is distinctive from that of the MT_1_-G_i_ complex.

### MT_1_ shares a similar G_s_ entry angle with GPR61 but shows a different binding mode from GPR61

To identify other GPCRs coupling to G_s_ in a similar way to MT_1_, we performed an amino acid sequence alignment of TM3 and ICL2 of class A GPCRs using the GPCRdb server ^25^. Surprisingly, only 5 class A GPCRs contain a Tyr at position 3.53, including MT_1_ and MT_2_ (Fig. 3a). Among the 5 GPCRs, GPR61 is the only receptor for which the structure of G_s_ complex is available in PDB. Structural comparison with the recent cryo-EM structure of the GPR61-G_s_ complex (PDB 8KGK) revealed that MT_1_ and GPR61 share a similar G_s_ entry angle (Fig. 3b, d). However, G_s_ enters deeper into GPR61 than into MT_1_ both at ICL2 and the cytoplasmic pocket of the receptor (Fig. 3c, e). At the ICL2 interface, residues around Tyr148^34.51^ of GPR61 make closer contacts than MT_1_, and Glu151^3.54^ of GPR61 interacts with G_s_ via Gln35^G.HN.52^, which i not observed in the MT_1_-minG_s_ (structure Fig. 3c). At the cytoplasmic pocket, the backbone of Tyr143^3.53^ of GPR61 forms a hydrogen bond with His387^G.H.5.19^ of G_s_, while Tyr128^3.53^ of MT_1_ forms a hydrogen bond with Gln390^G.H.5.22^ of G_s_ (Fig. 3e). Associated with this conformational difference, Pro147^34.50^ of GPR61 does not interact with G_s_, which is at the same position as Ser132^34.50^ of MT_1_ interacting with His387^G.H.5.19^ of G_s_. As a result, G_s_ enters deeper into GPR61 than MT_1_, associated with additional interactions; Gln347^8.49^ to Gln390^G.H.5.22^, and Arg346^8.48^ and Arg348^8.50^ to Glu392^G.H.5.24^. These structural insights may reflect the constitutive G_s_ signaling activity of GPR61 and the fact that G_s_ corresponds only to the secondary coupling of MT_1,_ which is primarly coulpled to G_i_.

**Fig. 3.**
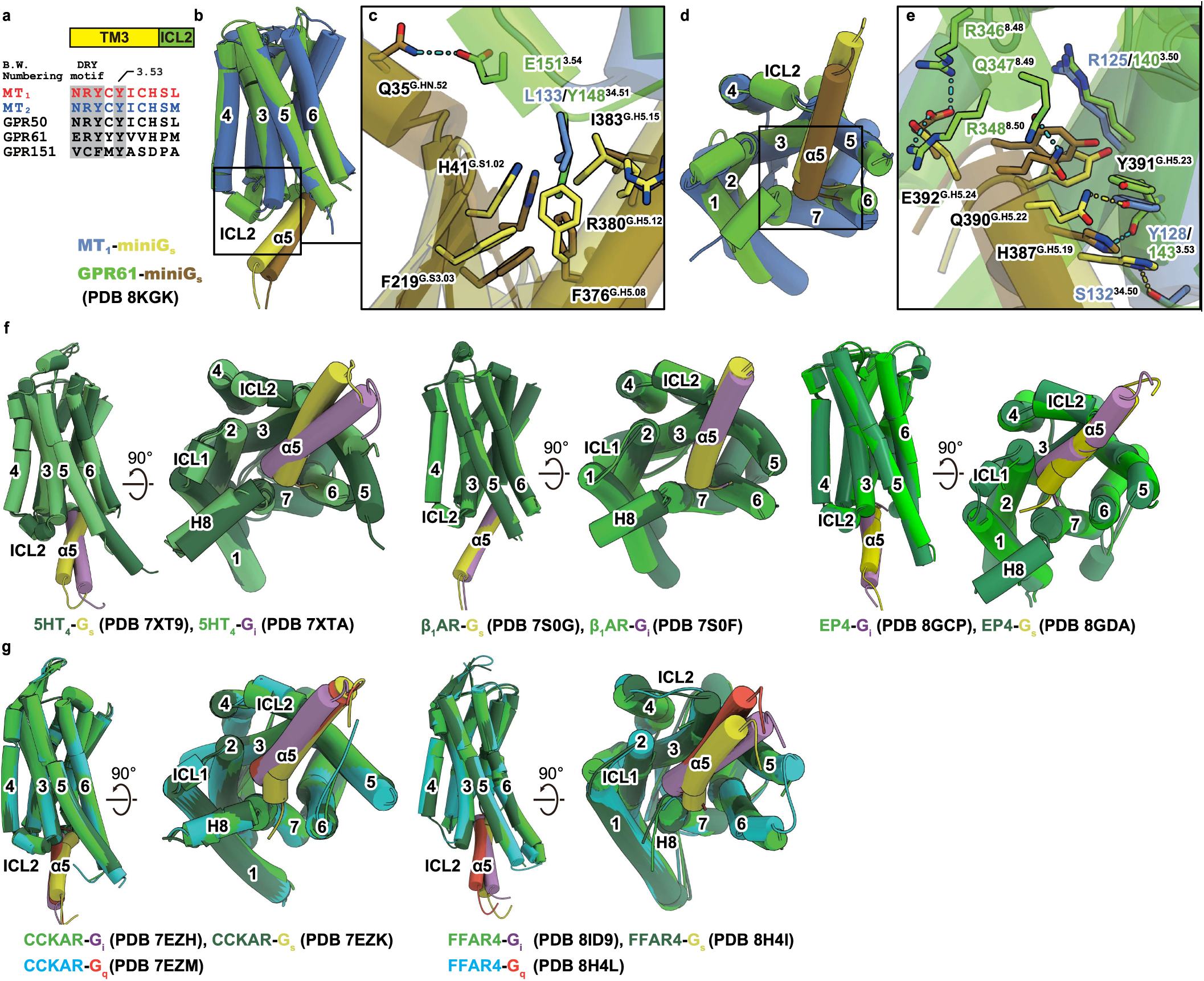
Structural comparison with GPR61-G_s_ complex and other G_s_/G_i_ complexes with the same receptor. (a), Amino acid sequences alignment of human MT_1_, MT_2_, GPR50, GPR61 and GPR151. (b-e), Structural comparison between MT_1_-miniG_s_ and GPR61-miniG_s_ (PDB 8KGK) displaying (b) overall, (c) ICL2 interface, (d) overall structure from cytoplasmic side and (e) α5 helix interface. (f, g), Structural comparison among reported GPCR-G complexes with the same receptor showing (f) primarily G_s_ coupling receptors including 5HT_4_ (PDB 7XT9 and 7XTA), β_1_AR (PDB 7S0G and 7S0F), EP4 (PDB 8GCP and 8GDA), or (g) primarily G_q_ coupling receptors including CCKAR (PDB 7EZM, 7EZH and 7EZK) and FFAR4 (PDB 8H4L, 8ID9 and 8H4I).

### Binding features among G_s_- and G_i_-signaling complexes

To understand how GPCRs couple to different G protein subtypes, we compared structures of five receptors with both G_s_-bound and G_i_-bound complexes: 5HT_4_ ^2^ (G_i_: PDB 7XTA, G_s_: PDB 7XT9), FFAR4 (G_i_: PDB 8ID9, G_s_: PDB 8H4I), β_1_AR ^3^ (G_i_: PDB 7S0F, G_s_: PDB 7S0G), CCKAR ^26^ (Gi: PDB 7EZH, G_s_: PDB 7EZK) and EP4 ^4^ (G_i_: PDB 8GCP, G_s_: PDB 8GDA). 5HT_4_, β_1_AR and EP4 primarily couple to G_s_, while CCKAR and FFAR4 primarily couple to G_q_. MT_1_ is unique as the first primarily G_i_i-coupled GPCR also complexed with G_s_. 5HT4-G_s_/-G_i_ and FFAR4-G_s_/-G_i_ displayed significant differences in the N-terminal region of α5 helix while maintaining similar C-terminal conformations, a pattern closely resembling the structural variations observed in MT_1_-G_s_/-G_i_ complexes. In contrast, EP4-G_s_/-Gi and CCKAR-G_s_/-Gi complexes exhibited more pronounced conformational differences at the C-terminal “wavy hook” region of α5. In β_1_AR complexes, G_s_ and G_i_ showed the closest overlay of α5, likely due to the engineered Gi protein containing N-terminal G_s_ substitutions. These structural comparisons reveal receptor-dependent mechanisms for dual G protein coupling with no universal structural determinant emerging from current structures, highlighting the complexity of GPCR-G protein selectivity.

### Structural basis of MT_1_ subtype specific G_s_-coupling is in the TM5-ICL3-TM6 region

Tyr128^3.53^ is a key feature of MT_1_ for G_s_ coupling, yet Tyr^3.53^ exists also in MT_2_, which does not couple to G_s_, suggesting that features other than Tyr^3.53^ explain the exclusive MT_1_ coupling to G_s_ and not MT_2_. Recent structural studies demonstrated that the distance from the latter half of TM5 to TM6 is the key for G_s_ and G_i_ selectivity of serotonin receptor subtypes, as swapping of the TM5-ICL3-TM6 domain between the G_s_-coupled 5HT_4_ and the G_i_-coupled 5HT_1A_ switched also their G protein coupling profiles^2^. Inspired by this study, we exchanged the TM5-ICL3-TM6 regions between MT_1_ and MT_2_ (Fig. 4a,b). The **MT_1_**-MT_2_(TM5-6) chimera lost the melatonin-induced cAMP elevation phase seen in wild-type MT_1_, whereas the **MT_2_**-MT_1_(TM5-6) chimera gained the cAMP elevation phase of the wild-type MT_1_ (Fig. 4c). The **MT_2_**-MT_1_(TM5-6) chimera coupled to both miniGα_i_ and miniGα_s_ proteins (Fig. 4d). These data indicate that the molecular determinants of G_s_ coupling are located in the TM5-ICL3-TM6 region of MT_1_ and are absent in MT_2_.

**Fig. 4.**
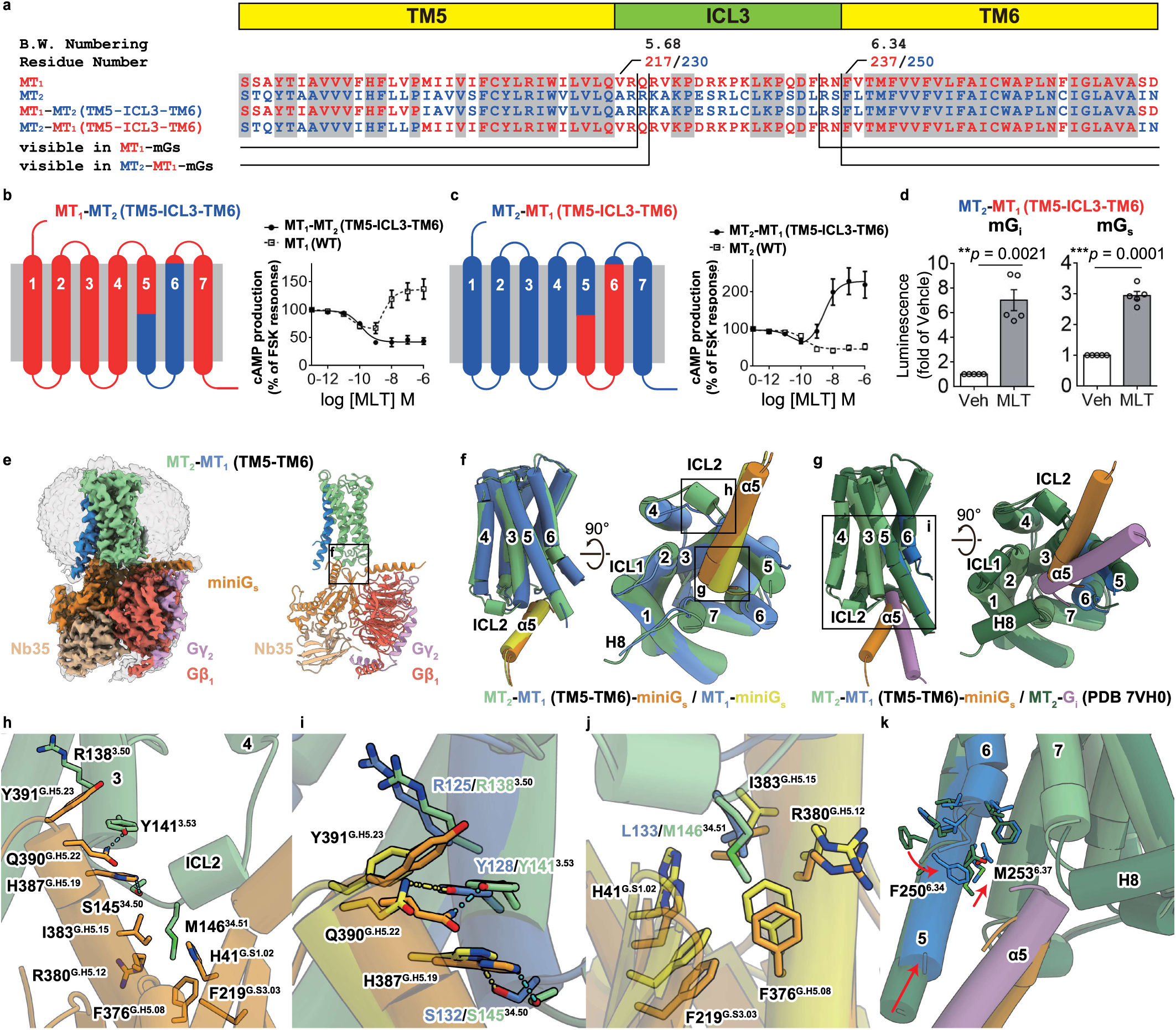
Structural basis of MT_2_-MT_1_-chimera Gs coupling. (a) Amino acid sequences alignment of human MT_1_ and MT_2_ chimeric receptors in TM5-ICL3-TM6 region. (b,c) Concentration–response effect of melatonin on FSK-induced cAMP production (5 min, FSK [5µM]) in HEK293T cells expressing (b) MT_1_–MT_2_(TM5– ICL3–TM6) or (c) MT_2_–MT_1_(TM5–ICL3–TM6) chimeric receptors, assessed with the BRET-based CAMYEL biosensor. Data represent means ± SEM of six independent experiments. (d) NanoBiT complementation assay showing recruitment of LgBiT–miniG_i_ (left) or LgBiT–miniG_s_ (right) to the MT_2_–MT_1_(TM5–ICL3–TM6)–NP receptor upon melatonin stimulation (1 µM, 2 min). Data present means ± SEM of five independent experiments. Statistical significance was determined using paired two-tailed t-tests, exact p-values are reported. (e) CryoEM density map (left) and overall structure (right) of MT_2_-MT_1_(TM5-TM6)-miniG_s_ with Gβ_1_, Gγ_2_ and Nb35. MT_2_, MT_1_(TM5-TM6), miniG_s_, Gβ_1_, Gγ_2_ and Nb35 were shown as light green, blue, orange, red, pink and brown, respectively. Map including micelle density was shown in transparent style. (f, g) Structural comparison of MT_2_-MT_1_(TM5-TM6)-miniG_s_ with (f) MT_1_-miniG_s_ or (g) MT_2_-G_i_ (PDB 7VH0), including lateral and cytoplasmic viewpoints. MT_2_ in MT_2_-G_i_ is shown as light purple, and G_i_ in MT_2_-G_i_ is shown as green. (h) The interface between MT_2_-MT_1_(TM5-TM6) and miniG_s_ around TM3 and ICL2. (i, j) Structural comparison between MT_2_-MT_1_(TM5-TM6)-miniG_s_ and MT_1_-miniG_s_ at (i) TM3 and α5 helix or (j) ICL2. (k) Structural comparison between MT_2_-MT_1_(TM5-TM6)-miniG_s_ and MT_2_-G_i_ at TM5, TM6 and α5 helix.

To better understand the binding mode of G_s_ to the **MT_2_**-MT_1_(TM5-6) chimera, we determined its cryo-EM structure in complex with miniG_s_ ^star 27^ (Fig. 4e). The overall structure of the **MT_2_**-MT_1_(TM5-6)-G_s_ complex was similar to the MT_1_-G_s_ complex (Fig. 4f), with the exception of the location of the α5 helix of the chimera complex which was slightly different from that of the MT_1_-G_s_ (Fig. 4f). Associated with this conformational change, Arg138^3.50^ formed a stacking interaction with Tyr391^G.H.5.23^ of G_s_, though Tyr141^3.53^ and Ser145^34.50^ kept interactions with Gln390^G.H.5.22^ and His387^G.H.5.19^ of G_s_, respectively (Fig. 4h,i). In parallel, Met146^34.51^ at the same position of Leu133^34.51^ maintained a hydrophobic interaction with the pocket of G_s_ (Fig. 4j). To understand why the substitution of TM5-ICL3-TM6 enabled MT_2_ to couple to G_s_, we compared the **MT_2_**-MT_1_(TM5-6)-G_s_ structure with the previously solved structure of MT_2_ in complex with G_i_ (Fig. 4g, k). Similar to MT_1_, different entry angles were observed between G_s_ and G_i_. No other substantial changes were found. The position of TM5 and TM6 was similar between MT_2_ and the **MT_2_**-MT_1_(TM5-6) chimera structures indicating that the positioning and the rotation of TM5 and TM6 is not responsible for G_s_ coupling of the chimera (Fig. 4k). Collectively, these data suggest that TM5 and TM6 do not play an important role in G_s_ coupling of MT_1_ vs MT_2_, and suggest that key molecular determinants are likely located within ICL3.

### ICL3 of MT_1_ supports coupling to G_s_ but not to G_i_

To provide experimental evidence for the importance of ICL3 over TM5 and TM6 of MT_1_ for G_s_ coupling, we generated MT_2_-MT_1_ chimera containing TM5, ICL3 or TM6 of MT_1_ alone or a combination of ICL3 and TM6 (Fig. 5a-e, Supplementary Fig.S3). All chimeras were expressed at comparable levels as the MT_2_-MT_1_(TM5c-ICL3-TM6) reference chimera (Supplementary Fig. S4). All chimeras showed the melatonin-induced cAMP inhibition phase (Fig. 5b-e, Supplementary Fig.S3). The MT_2_-MT_1_(ICL3) chimera displayed, in addition, the cAMP elevation phase (Fig. 5d). Among the 20 amino acids of the ICL3 region, the MT_2_-MT_1_(coreICL3) chimera containing the N-terminal 12 amino acids (VRQRVKPDRKPK) of MT_1_ maintained the G_s_ phase (Fig. 5e), whereas the MT_2_-MT_1_(C-ICL3+TM6) chimera containing the 8 C-terminal amino acids did not contribute to the G_s_ phase (Supplementary Fig. S3b). Further delimitation of the 12 amino acid stretch failed to induce the G_s_ phase in the MT_2_ context (Supplementary Fig. S3c,d), defining the 12 N-terminal amino acid of the MT_1_-ICL3 as the minimal region for the G_s_ phase of cAMP signaling. Coupling of the minimal MT_2_-MT_1_(coreICL3) chimera to both G_s_ and G_i_ was confirmed directly in the miniG_s/i_ coupling assays (Fig. 5f,g).

**Fig. 5.**
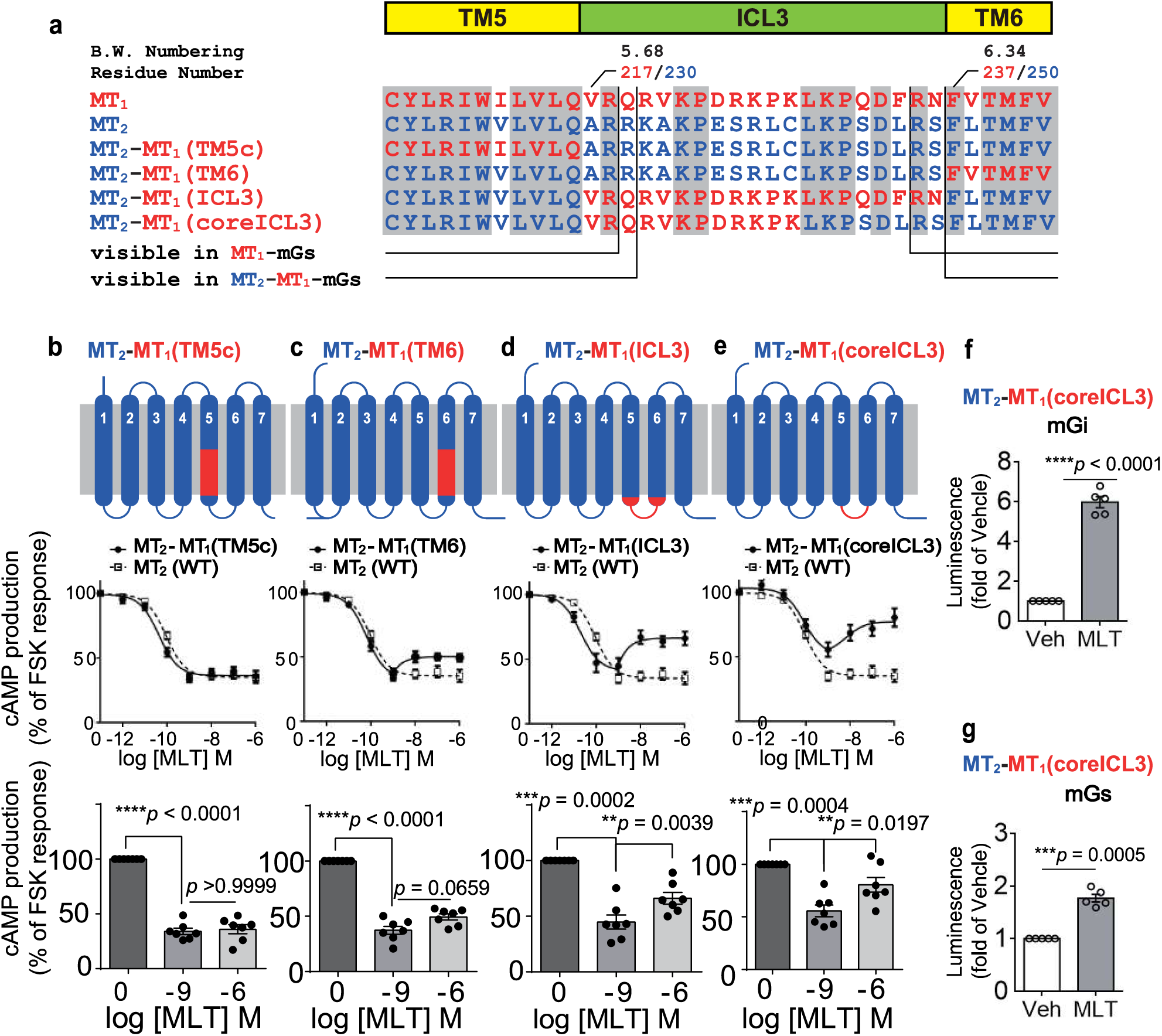
Identification of key determinants of MT_1_-G_s_ interaction by MT_2_-MT_1_ chimera receptors. (a) Amino acid sequences alignment of human MT_2_ chimeric receptors. (b-e) (upper) Schematic representation of chimera. (middle) Concentration– response effect of melatonin on FSK-induced cAMP production (10 min, FSK [5µM]) in HEK293T cells expressing (b) MT2–MT1(TM5c), (c) MT2–MT1(TM6), (d) MT2–MT1(ICL3) or (e) MT2–MT1(core ICL3) chimeric receptors, assessed with the BRET-based CAMYEL biosensor. Data represent means ± SEM of seven independent experiments. (lower) Statistical comparison of concentration–response data at 0, 1 nM and 1 µM of melatonin. Data represent means ± SEM of six independent experiments. Statistical analysis was performed by one-way ANOVA followed by Bonferroni’s multiple comparison test. (f, g) NanoBiT complementation assay showing recruitment of LgBiT–miniGi (f) or LgBiT–miniGs (g) to the MT_2_–MT_1_(coreICL3)–NP receptor upon melatonin stimulation (1 µM, 2 min). Data represent means ± SEM of five independent experiments. Statistical significance was determined using paired two-tailed t-tests, exact p-values are reported.

Since the flexible ICL3 could not be resolved in our MT_1_-G_s_ structure, we performed molecular dynamics (MD) simulations of MT_1_-G_s_, MT_1_-G_i_, MT_2_-G_i_, and MT_2_-MT_1_(TM5-ICL3-TM6)-G_s_ complexes using AlphaFold3-modeled ICL3 structures. A MT_2_-G_s_ test model was generated to understand why wild-type MT_2_ cannot stabilize G_s_ interactions. Unlike many GPCRs with extensive ICL3 regions (>100 residues), the melatonin receptors possess short, 20-residue, loops with high AlphaFold3 confidence (>60%). MD simulations revealed that the ICL3 forms dynamic polar and hydrophobic interactions with G protein α5 helices, with notable hydrophobic contacts between MT_1_ V217^ICL3^ and L388^G.H5.20^ of G_s_ (Fig. 6a,b, Extended Data Fig. 6a). Together with I129^3.54^, V214^5.65^, and L213^5.64^, these residues create a hydrophobic patch that stabilizes the receptor-G_s_ interface. While V217^ICL3^ also interacts with G_i_, these contacts are less essential because Gi α5 penetrates deeper into TM5/TM6 regions, achieving stability through intrinsic hydrophobic interactions involving L348^G.H5.20^, F354^GH5.26^, and L353^GH5.25^ with I210^5.61^, whose alanine mutation substantially reduces G_i_ coupling ^12^. The predicted importance of V217^ICL3^ for G_i_ and G_s_ coupling was tested in the context of the MT_2_-MT_1_(coreICL3) chimera representing the minimal MT_1_ sequence enabling notable MT_2_-G_s_ coupling, by the V217A^ICL3^ mutant replacing Val 217 by Ala present in MT_2_ (Fig. 6c). While the inhibitory phase was preserved, the stimulatory phase was abolished consistent with the important role of V217^ICL3^ for G_s_, but not G_i_, coupling (Fig. 6c). Deletion of ICL3 (ΔQRVKPDRKPK) in MT_1_ resulted in the loss of the melatonin-induced cAMP stimulation phase while maintaining the inhibition phase further confirming the essential role of ICL3 for G_s_ coupling but not for G_i_ coupling to MT_1_ (Fig. 6d). This ICL3 deletion chimera also shows that V217^ICL3^ alone is not sufficient for G_s_ coupling as this residue as well as the other residues involved in the hydrophobic patch (I129^3.54^, L213^5.64^, V214^5.65^) are present in this chimera (Fig. 6b). This is consistent with the observation that chimeric replacement of TM5 alone does not confer G_s_ coupling to MT_2_ (see Fig. 5b). Our MT_2_-G_s_ test model MD simulations support this mechanism, showing that equivalent alanine residues in MT_2_ ICL3 cannot provide adequate stabilization of G_s_, resulting in high ICL3 and α5 mobility with substantially reduced interaction energies (Extended Data Fig. 6b-d). Since VRQRVKPDRKPK represents the minimal sequence enabling notable MT_2_-G_s_ coupling, specific ICL3 conformation appears also essential for establishing hydrophobic interactions necessary for G_s_ recognition. Our findings demonstrate that ICL3 is essential for G_s_ coupling but dispensable for G_i_ coupling, in a context where G_s_ coupling to MT_1_ relies heavily on ICL3 interactions, which likely compensates for its shallow binding mode.

**Fig. 6.**
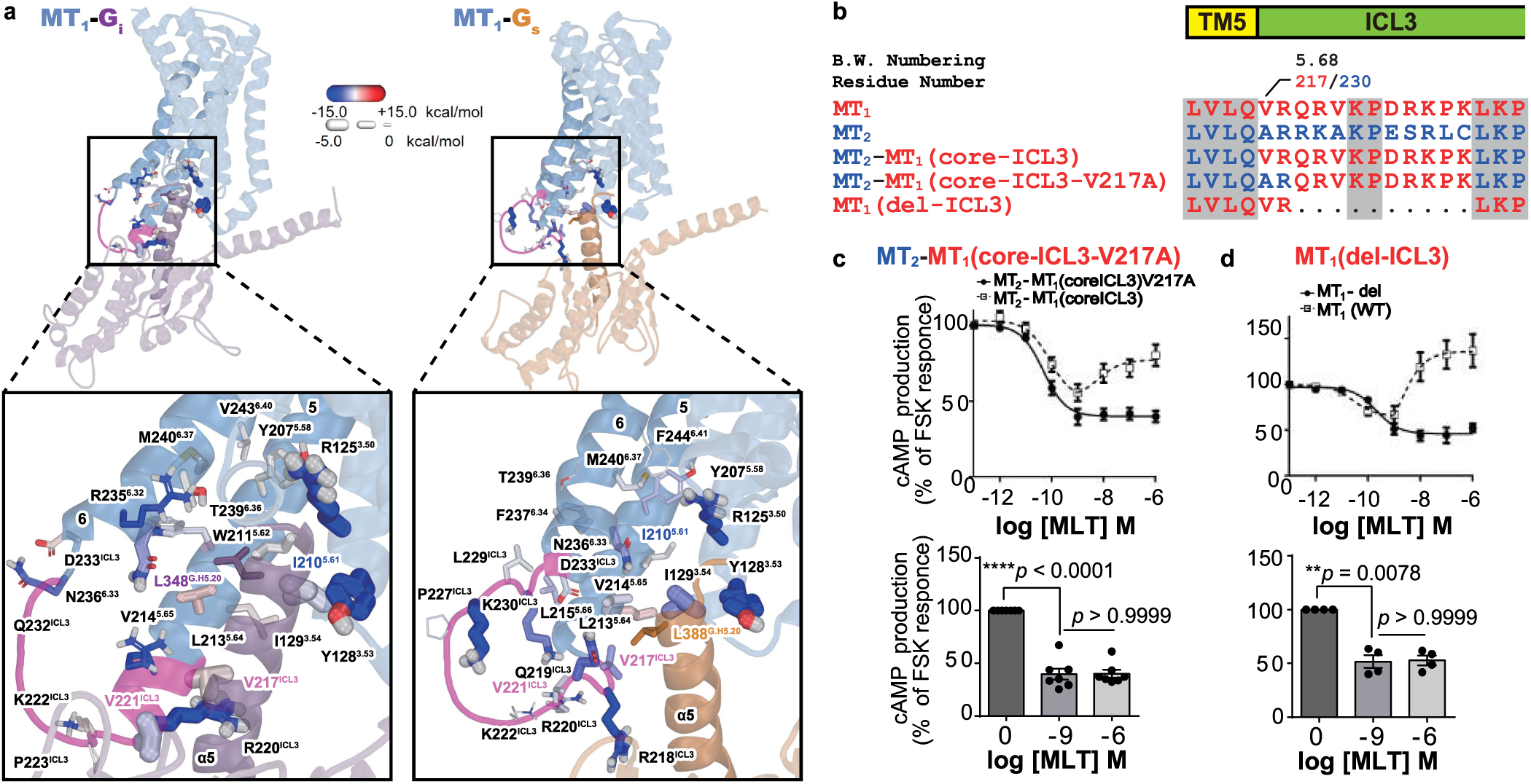
Molecular dynamics simulations support differential G protein binding modes and ICL3-dependent Gs coupling of MT1. (a) Representative MD simulation snapshots of MT_1_-G protein complexes. Carbon colours indicate electrostatic energies, stick thickness represents van der Waals interactions with α5 helix. Only residues >1 kcal/mol in TM5-ICL3-TM6 are shown. AlphaFold3-modeled ICL3 (pink) highlights non-conserved hydrophobic residues. TM5 and TM6 are highlighted in darker colour. Key contacts: I210^5.61^ with L348^G.H5.20^, F354^GH5.26^, and L353^GH5.25^ (Gi deep binding) versus V217^ICL3^ with L388^G.H5.20^ (Gs shallow binding). (b) Amino acid sequence alignment of human MT_2_ chimera and human MT_1_ deletion mutant in TM5-ICL3 region. (c,d) (upper) Concentration–response effect of melatonin on FSK-induced cAMP production (10 min, FSK [5µM]) in HEK293T cells expressing (c) MT2-MT1(coreICL3-V217A) and (d) MT1-ICL3-deletion mutant chimeric receptors, assessed with the BRET-based CAMYEL biosensor. Data represent means ± SEM of seven (c) or four (d) independent experiments. (Lower) Statistical comparison of concentration–response data (Upper) at 0, 1 nM and 1 µM of melatonin. Data represent means ± SEM of six independent experiments. Statistical analysis was performed by one-way ANOVA followed by Bonferroni’s multiple comparison test.

## Discussion/conclusion

We describe here the molecular determinants enabling G_s_ coupling to MT_1_, a prototypical G_i_-coupled GPCR. The binding mode of the MT_1_-G_s_ complex is structurally distinctive from the MT_1_-G_i_ coupling: while G_i_ binding involves the C-terminal end of TM5 and TM6 of MT_1_^12,13^, G_s_ forms essential interactions with TM3 (Y128^3.53^) and ICL2 (L133^34.51^) of MT1, which were experimentally validated. Since the flexible ICL3 could not be resolved in our MT_1_-G_s_ structure, we investigated its role through MT_2_-MT_1_ chimera studies and MD simulations. Chimera studies defined the N-terminal 12 amino acids (VRQRVKPDRKPK) of MT_1_-ICL3 as minimal region required for G_s_ coupling with V217^ICL3^ establishing hydrophobic contacts participating in G_s_ recognition, stabilizing the otherwise shallow binding mode of G_s_ in terms of depth of penetration into the TM5/TM6 region. Consistently with the absence of G_s_ coupling to MT_2_, the stabilizing role of ICL3 is not seen for this receptor in MD simulations. While the ICL3 is essential for G_s_ coupling to MT_1_, it is dispensable for G_i_ coupling as demonstrated with the MT_1_-ICL3 deletion mutant.

The contribution of ICL3 in G protein coupling has been suggested previously by functional and bioinformatics studies^28–30^. On the other hand, for different serotonin receptor subtypes, the length of TM5 and TM6 has been suggested as important determinant of G_s_-G_i_ selectivity^2^. For MT_1_, the contribution of TM5 and TM6 could be excluded by solving the cryo-EM structure of the **MT_2_**-MT_1_(TM5-6) chimera which couples to G_s_, but shows no significant differences in TM5 and TM6 length, position and rotation compared to the MT_2_, which does not couple to G_s_. The study of Sadler et al. ^30^ suggested a gating function of ICL3 controlling G access to receptors. Our data suggest different roles of ICL3 for G protein coupling to MT1. Whereas the MT_1_-G_s_ complex suggests rather a supportive role of ICL3 for G_s_ coupling, ICL3 is dispensable for G_i_ coupling to MT_1_.

Coupling of MT_1_ to G_i_ occurs upon acute melatonin stimulation. Coupling to G_s_ is also of physiological relevance as it occurs upon long-term melatonin stimulation, mimicking the conditions at dawn, when receptors are exposed to melatonin throughout the night. Stimulation of the G_s_/cAMP/PKA pathway has been observed in our study *ex vivo* and *in vivo* in the mouse PT expressing endogenous MT_1_. Previously, melatonin has been shown to have a dual impact on the rhythmic transcription of PT cells ^31^. In the beginning of the night melatonin acutely suppresses cAMP levels which is consistent with the well-documented G_i_-coupling of MT_1_ receptor. At dawn, melatonin has a delayed heterologous sensitizing effect on the adenylyl cyclase system^32^ which was shown by the sensitization of the signaling of adenosine A_2b_ receptor, the most prominently expressed G_s_-coupled receptor in the PT^33^. Our study indicates that apart from this heterologous sensitization by amplifying the A_2b_ receptor response, the G_i_-G_s_ switch of MT_1_ coupling is likely to contribute to the increased activity of the cAMP/PKA pathway at the end of the night, thus enhancing the contrast of cAMP-dependent rhythmic transcription in PT cells at the night/day transition. These results provide new insights on the molecular mechanisms underlying the synchronization effects of melatonin, where the acute G_i/o_ pathway upon melatonin rise would signal the beginning of the dark phase of the light/dark cycle, and the switch to G_s_-coupling would signal the night/day transition.

## Materials and Methods

### Material

HEK293T cells were used as described previously^34^. The expression constructs of melatonin receptors used in this study were as follows. For Fig. 1a,b, C-terminal untagged Flag-MT_1_ and HA-MT_2_ constructs were used as described previously^34^. For other experiments, C-terminal–tagged constructs (Flag-MT_1_-NP and Flag-MT_2_-NP) were generated by PCR-based modification using plasmids obtained from Addgene (Plasmid #66443^35^ and #66444^35^). Melatonin receptors (MTR) mutants were generated by PCR using PNK-phosphorylated primers carrying the intended mutation, followed by self-ligation with T4 DNA ligase (Nippon Gene), as described previously^36^. Chimeric MTRs were generated by PCR using the In-Fusion HD cloning kit (Clontech) as described previously^37^. cDNA encoding LgBit-miniGs protein or –miniGs/i chimera protein was provided by Dr. Nevin Lambert (Augusta, Georgia, USA).

### Cell Culture and Transfection

HEK293T cells were transiently transfected with Jet-PEI or PEI-Max according to the manufacturer’s protocol^36,37^. Briefly, approximately 0.3 × 10⁶ cells per well were seeded into 12-well plates. After 6 h, a total of 1 µg DNA (expression plasmid supplemented with empty vector) was mixed with 3 µL of transfection reagent, incubated for 15 min, and then added to the cells. On the following day, cells were transferred to white 96-well plates, and experiments were performed 48 h post-transfection.

### miniG protein Nanoluc Complementation assay

Nanobit complementation assay was performed as following: HEK293T cells seeded in 12 well plate were co-transfected with LgBiT-fused miniGs (5ng/well) or miniGs/i (15ng/well) together with 5ng/well NP-fragment–tagged^21^ human Flag-MT_1_ or Flag-MT_2_ receptors or their chimera proteins. Cells were reseeded into white 96-well plates on the following day. After 46 h, the culture medium was replaced with phenol red–free DMEM. Two hours later, 5 µM coelenterazine H was added, and baseline luminescence was recorded. Cells were then stimulated with vehicle or 1 µM melatonin, and luciferase activity was measured. Data represent luminescence values obtained 2 min after melatonin stimulation.

### *In Vitro* cAMP Assay by HTRF

cAMP accumulation was assessed by HTRF (Fig.1a,b) under suspension conditions as described previously^34^. HEK293T cells were transfected with varying amounts of Flag-MT_1_ without any C-terminal tag (1 µg, “high”; or 10 ng plus 990 ng empty vector, “low”) or 1 µg HA-MT_2_ without any C-terminal tag plasmids in 12-well plates. After 48 h, cells were dispensed into 384-well plates (4,000 cells per well) and stimulated with 2 µM forskolin in the presence of the indicated melatonin concentrations for 30 min at room temperature in PBS supplemented with 1 mM 3-isobutyl-1-methylxanthine (IBMX; Sigma-Aldrich, France). Cells were then lysed, and cAMP levels were quantified following the manufacturer’s instructions (Cisbio Bioassays, France). Where indicated, cells were pretreated with PTX (200 ng/mL, 4 h). Luminescence was recorded using an Infinite F500 microplate reader (Tecan).

### cAMP monitoring by CAMYEL-BRET sensor

cAMP measurement by CAMYEL-BRET sensor was performed as previously described^38^. HEK293T cells were co-transfected with 500 ng CAMYEL sensor and 500 ng melatonin receptor (Flag-MT_1_-NP or Flag-MT_2_-NP and their mutants or chimera receptor) plasmids in 12-well plates. The following day, cells were reseeded into 96-well plates. After 46 h, the culture medium was replaced with phenol red–free DMEM. Two hours later, 5 µM coelenterazine H was added, and baseline BRET signals were recorded. Cells were then treated with vehicle or 5 µM forskolin in the presence of the indicated melatonin concentrations (without IBMX), and real-time BRET changes were monitored. Data shown correspond to measurements taken 10 min after melatonin stimulation. BRET was measured by TECAN SPARK.

### Animals

Five-week-old male C57BL/6JJmsSlc mice (Japan SLC Inc., Shizuoka, Japan) were purchased and housed in plastic cages (170 W × 240 D × 125 H mm, Clea, Tokyo, Japan) under a 12-h light (200 lx of fluorescent light) / 12-h dark cycles (12L12D, 0800 light ON, 2000 light OFF), maintained at a constant temperature (23 ± 1 °C). Food (CE-2; CLEA) and water were provided *ad libitum*. All animal experiments were approved by the Committee of Animal Care and Use of the Aichi Medical University. All experimental procedures were conducted in accordance with the institutional guidelines for the use of experimental animals.

### *Ex-* and *In-Vivo* Melatonin treatment

For analysis of melatonin effect on cultured PT/MBH, brain of 10─12-week-old mice were quickly removed under general anesthesia at 6 hours after dawn and immersed in ice-cold Hank’s buffered saline (HBS; #09735-75, Nacalai tesque, Kyoto, Japan). The hypothalamus, containing the PT and MBH was sectioned at 1 mm thickness by brain matrix. Then, the PT/MBHs were placed on a culture membrane (Millicell-CM PICM0RG50, Merck Millipore, Darmstadt, Germany) in a 35-mm petri dish and cultured at 37°C with 1.2 mL of medium containing Dulbecco’s modified Eagle medium (DMEM) (Gibco, Carlsbad, CA), 100 U/mL penicillin, and 100 μg/mL streptomycin. After 6 hours preincubation, slices were then added with the culture medium containing melatonin at ZT12 (finally 10 μM, 0.1 % DMSO; Wako, Tokyo, Japan). After 16 hours, proteins of the PT/MBH were extracted for using cAMP measurement and western blotting.

For *in vivo* analysis of melatonin effect, mice were kept under 8-h light / 16-h dark cycles (8L16D) for 3 weeks, and melatonin (0.26 mM / 0.1mL) were intraperitoneally (i.p.) injected as previous report 1, once at ZT8. After 16 hours (ZT0), mice were perfused then fixed on slides with 4% paraformaldehyde (PFA) in 0.1 M phosphate buffer (pH 7.4), and brains were removed and put into the 4% PFA for 48 hours, and then transfer to 20 % sucrose / PBS for using frozen sectioning of brain.

### *Ex vivo* cAMP measurement at PT

Protein extraction of cultured PT/MBH was performed using cell lysis buffer (#9803, Cell Signaling Technology, Tokyo, Japan) containing protease inhibitor cocktail (#P8340, Sigma) and phosphatase inhibitor cocktail 1 (#P2850, Sigma) according to the manufacturer’s instructions.

cAMP measurements were performed with a homogeneous TR-FRET immunoassay using the LANCE cAMP Detection Kit (#AD0262, PerkinElmer, USA), according to the manufacturer’s instructions (PerkinElmer). Cultured PT/MBH tissues homogenates (0.2 μg/μL) were diluted 10 times with stimulation buffer, and added to 10 µL of Alexa Fluor 647 anti-cAMP antibody diluted with stimulation buffer. After incubation for 45 min at room temperature, the reaction was stopped by the addition of 20 µL working solution (10 µL Eu-cAMP and 10 µL ULight-anti-cAMP), and incubated for 1 h at room temperature. The TR-FRET signal was read using a microplate reader SpectraMax M5 (Molecular Devices). cAMP concentrations were determined using GraphPad Prism 8 software (GraphPad Software Inc., San Diego, CA).

### Western blot analysis

Western blot analysis was performed as follows: homogenized PT/MBH was loaded as 2 μg protein samples per lane onto 8 % SDS-polyacrylamide gels. Following separation at 100 mA, membranes were incubated with the following primary antibodies: rabbit monoclonal antibodies against phospho-CREB (Ser133) (1:1000; #9198, Cell Signaling Technology), and CREB (1:1000; #9192, Cell Signaling Technology). Membranes were washed and then incubated with HRP-conjugated goat polyclonal antibody against rabbit IgG (1:10,000; #7074, Cell Signaling Technology). By using ImmunoStar LD (#292-69903, Fijifilm), chemiluminescent images were detected using an Amersham Imager 600 (Cytiva Lifescience).

### Immunohistochemistry

Immunohistochemistry was performed by using the Vectastain Elite ABC rabbit IgG kit (Vector, Burlingame, CA) as our previous reports^39^. Frozen sections of paraformaldehyde-fixed trimmed mice brain (14 µm) including the PT/MBH were used. Briefly, after drying, the sections were immersed in HistoVT One (pH7.0, #06380, Nakarai) and heated in a microwave oven for 15 min at low voltage. After blocking with normal rabbit serum (Vectastain), sections were incubated with rabbit polyclonal antibody against phospho-CREB (Ser133) (1:500; #9198, Cell Signaling Technology) at 4°C for 48 h. After encapsulation with mounting medium, images were detected using a BZ-X800 (Keyence). ImageJ software (NIH) was used to analyze pCREB-immunoreactive (ir) cell numbers as previous report^40^. Data are shown as percent of maximum values (pCREB-ir cells /mm2).

### Expression and purification of Gβ_1_-Gγ_2_ dimer

In the previous study, we expressed and purified the Gi1 heterotrimer including human Gαi1, bovine Gβ_1_ and mouse Gγ_2_ by Ni-NTA affinity chromatography and anion exchange chromatography. As a byproduct of the anion exchange chromatography step, the fraction of Gβ_1_-Gγ_2_ dimer was eluted before the fraction of the Gi_1_ heterotrimer.

### Expression and purification of Nb35

The plasmid encoding Nb35 was prepared as previously reported^41^. The protein was expressed in the periplasm of E. coli C41 (Rosetta) cells cultured in LB medium supplemented with 1 mM IPTG for 20 h at 25 °C. After 20 h, the cells were collected and disrupted by ultrasonication in hypotonic buffer (20 mM Tris-HCl, pH 7.5, 150 mM NaCl and 2 mM MgCl2), and the Nb35 protein was purified by Ni-NTA affinity chromatography and then subjected to size-exclusion chromatography on a HiLoad Superdex75 16/600 column. Peak fractions were pooled and concentrated to 3 mg ml^−1^.

### Expression and purification of MT_1_-miniG_s_-Gβ_1_-Gγ_2_ complex

The HA signal sequence, the FLAG epitope tag (DYKDDDDK), the ALFA tag (SRLEEELRRRLTE)^42^ and GSGSG linker were fused to the N-terminus of MT_1_, in this order. In addition, we fused the miniGs fragment to the C-terminus of MT_1_.

In the MT_2_-MT_1_ chimera construction, we substituted TM5-ICL3-TM6 of MT_2_ to the MT_1_ sequence (from Met200^5.51^ to Phe256^6.53^) between Pro212^5.50^ and Ile270^6.54^. And we fused the same fragments as MT_1_-G_s_ to the N-terminus of MT_1_. In contrast, we fused mini-Gs^star^ protein created in our previous study^41^ to the C-terminus of MT_1_. All the above constructions were cloned into the pEGBacmam vector.

Then we infected a one tenth volume of a solution containing the virus encoding the above construction to the HEK293F cells at a density of 3–4 × 10^6^ cells ml^−1^, and grown at 37 °C for 20 h in FreeStyle 293 Expression Medium (Gibco). After sodium butyrate (FujiFilm Wako Pure Chemical Corporation) was added to 10 mM, the cells were incubated further at 30 °C for 48 h.

The HEK293F cells were collected by centrifugation at 5,000g for 10 min, lysed in buffer containing 50 mM Tris (pH 8.0), 150 mM NaCl and 10% glycerol and fresh frozen by liquid nitrogen. The cells were thawed and directly solubilized at 4 °C for 2 h in the solubilization buffer, containing 50 mM Tris (pH 8.0), 150 mM NaCl, 10% glycerol,1.5% DDM, 0.15% CHS, 5.2 μg ml^−1^ aprotinin, 2.0 μg ml^−1^ leupeptin, 1.4 μg ml^−1^ pepstatin A, 100 μM PMSF and 10 μM melatonin (Wako). The soluble fraction containing MT_1_-miniG_s_ was separated by ultracentrifugation (186,000g for 30 min), and the supernatant was mixed with the purified Gβ_1_-Gγ_2_ dimer (0.8 mg l−1 HEK cells), the purified Nb35 (0.5 mg l−1 HEK cells) and final 50 mU/mL of Apyrase (New England Biolabs). Then they were stirred at 4 °C for overnight. The solution was supplemented with 2 mM CaCl_2_ at the final concentration, mixed with the M1 anti-FLAG resin and rotated at 4 °C for 2 h.

The resin was collected by centrifugation (500g for 3 min) and washed with 20 column volumes of buffer, containing 50 mM Tris-HCl (pH 8.0), 150 mM NaCl, 10% glycerol, 0.03% LMNG, 0.003% CHS, 2 mM CaCl_2_ and 1 μM melatonin. The complex was eluted with buffer, containing 50 mM Tris-HCl (pH 8.0), 150 mM NaCl, 10% glycerol, 0.03% LMNG, 0.003% CHS, 2 mM CaCl_2_, 1 μM melatonin, 5 mM EDTA (pH 8.0) and 125 μg ml^−1^ FLAG peptide (Synthesized by GenScript), concentrated and purified by size-exclusion chromatography on a Superdex200 10/300 Increase (GE) column, using buffer containing 50 mM HEPES-NaOH (pH 7.5), 150 mM NaCl, 100 μM TCEP, 0.01% LMNG, 0.001%\ CHS and 1 μM melatonin. Peak fractions were pooled and concentrated to around 7 mg ml^−1^ with a centrifugal filter device (Millipore 50 kDa molecular weight cutoff).

### Grid preparation and cryo-EM data collection

The purified complex solution was applied to freshly glow-discharged Au 300 mesh R1.2/1.3 grids (Quantifoil), using a Vitrobot Mark IV (FEI) at 4 °C, with a blotting time of 4 s under 100% humidity conditions. The grids were then plunge-frozen in liquid ethane cooled to the temperature of liquid nitrogen.

Cryo-EM data were collected using a Titan Krios G3i microscope (Thermo Fisher Scientific), running at 300 kV and equipped with a Gatan Quantum-LS Energy Filter (GIF) and a Gatan K3 Summit direct electron detector in the electron counting mode (The University of Tokyo, Japan). Movies were recorded at a nominal magnification of 105,000×, corresponding to a calibrated pixel size of 0.83Å, with a total dose of approximately 50 electrons per Å^2^ per 48 frames. The data were automatically acquired using the EPU software (Thermo Fisher Scientific), with a defocus range of −0.8 to −1.6 μm. For MT1-miniGs, 10,373 movies were obtained. For MT1-MT2-miniGs, 8,174 movies were obtained.

### Image processing

All acquired movies were dose-fractionated and subjected to beam-induced motion correction implemented in RELION 3.1^43^. The contrast transfer function (CTF) parameters were estimated using patch CTF estimation in cryoSPARC v3.3^44^.

For MT_1_-miniG_s_, particles were initially picked from a small fraction with the Blob picker and subjected to several rounds of two-dimensional (2D) classification in cryoSPARC using 897 movies. Selected particles were used for template picking on the full dataset, and 8,196,012 particles were picked and extracted with a pixel size of 3.32 Å, followed by 2D classification, ab initio reconstruction, homogeneous refinement and heterogeneous refinement, displayed in the Extended Data Fig. 1. A total of 624,555 particles were used for training of topaz model^45^, and 3,524,939 particles were extracted. After 2D classification, heterogenous refinement and non uniform (NU) refinement, 449,204 particles were transferred into RELION 3.1 and were processed by 3D classification without alignment using TMD mask with several T parameters. After 3D refinment with SIDESPLITTER^46^, particle polishing, re-extract as 1.0 Å/pix and removal of the duplicated particles, 85,855 particles were transferred into cryoSPARC.

Finally, these 85,855 particles were reconstructed using NU refinement, resulting in a 3.03 Å overall resolution reconstruction, with the gold standard Fourier Shell Correlation (FSC = 0.143) criteria in cryoSPARC. Moreover, the 3D model was refined with a mask on the receptor by local refinement. As a result, the local resolution of the receptor portion was estimated as 3.24 Å by cryoSPARC. In parallel, the 3D model was refined with a mask on the G protein by local refinement, resulting in the local resolution at 2.9 Å estimated by cryoSPARC.

For MT_2_-MT_1_-miniG_s_, particles were initially picked from a small fraction with the Blob picker and subjected to several rounds of 2D classification in cryoSPARC using 2,429 movies. Selected particles were used for template picking on the full dataset, and 54,229,298 particles were picked and extracted with a pixel size of 3.32 Å, followed by 2D classification, ab initio reconstruction, homogeneous refinement and heterogeneous refinement, displayed in the Extended Data Fig. 2. A total of 272,585 particles were selected as the best class and were transferred into RELION 4.0. After 3D classification without alignment using TMD mask with several T parameters, 74,839 particles were selected, processed by 3D refinement with SIDESPLITTER^46^, particle polishing and re-extract as 1.0375 Å/pix and transferred into cryoSPARC.

Finally, these 74,839 particles were reconstructed using NU refinement, resulting in a 2.94 Å overall resolution reconstruction, with the gold standard Fourier Shell Correlation (FSC = 0.143) criteria in cryoSPARC. Moreover, the 3D model was refined with a mask on the receptor by local refinement. As a result, the local resolution of the receptor portion was estimated as 3.43 Å by cryoSPARC. In parallel, the 3D model was refined with a mask on the G protein by local refinement, resulting in the local resolution at 2.82 Å estimated by cryoSPARC.

### Model building and validation

The cryo-EM structures of the MT_1_-Gi complex (PDB 7DB6)^12^, MT_2_-Gi complex (PDB 7VH0)^13^ and OR51E2-miniGs (PDB 8F76)^47^ were used as starting templates for modeling the MT_1_, MT_2_-MT_1_, miniGs, Gβγ and Nb35 components. Initially, these models were positioned into the density map using jiggle fit in COOT^48^. Then these models were modified by COOT and refined with phenix.real_space_refine (v1.19)^49,50^ and the secondary structure restraints from phenix.secondary_structure_restraints. Cif format definition (ML1) downloaded from PDB was used for modeling melatonin molecule at ligand binding site.

### Computer Simulations

MD simulations were conducted on melatonin receptor systems derived from cryo-EM structures. All simulated systems and their stability parameters are listed in **(Supplementary Table 1**). MD simulations were performed on five distinct melatonin receptor-G protein complexes: MT_1_+Gi (PDB: 7DB6), MT_1_+Gs, MT_2_+Gi (PDB: 7VH0), MT_2__MT_1_(TM5-ICL3-TM6)+Gs, and MT_2__WT+Gs. The cryo-EM structures were prepared for simulations using Schrödinger Maestro (2025-1)^51^. ICL3 regions were rebuilt by grafting the corresponding segments from GPCRdb active state AlphaFold2-Multistate models ^52,53^ (version 15-05-2024): MT_1_ AlphaFold ICL3 (V217-P231) with an average pLDDT confidence score of 65.17, and MT_2_ AlphaFold ICL3 (A230-P244) with an average pLDDT confidence score of 73.62. The grafting boundaries were determined by identifying structured TM5/TM6 regions flanking the missing ICL3 density in the cryo-EM structures. AlphaFold segments were integrated as follows: MT_1_+Gi [V198-P244], MT_1_+Gs [V198-N236], MT_2_+Gi [L211-L251], MT_2__MT_1_(TM5-ICL3-TM6)+Gs [V198-P254]. The experimentally determined chimeric structure was used as the template for both MT_2__MT_1_(TM5-ICL3-TM6)+Gs and MT_2__WT+Gs systems, with the MT_2__WT+Gs variant generated by mutating the MT_1_-derived TM5-ICL3-TM6 segment back to MT_2_ wild-type sequence using Schrödinger Maestro 3D builder utility. In all systems, D55^2.50^ was protonated as per PROPKA3 predictions^54,55^. CHARMM-GUI^56–64^ server was used to prepare simulation systems. Membrane protein orientations were determined using the Positioning of Proteins in Membranes (PPM) 2.0^56^ web server prior to embedding in a 1-palmitoyl-2-oleoyl-sn-glycero-3-phosphocholine (POPC) bilayer membrane. Membrane dimensions were optimized for each receptor-G protein complex, ranging from 130 Å × 130 Å to 140 Å × 140 Å, to maintain adequate lipid-protein boundaries and prevent periodic artifacts. Each system was solvated with TIP3P water extending 30-35 Å from the membrane surfaces, with ionic strength adjusted to physiological conditions using 0.15 M NaCl. The resulting simulation systems comprised approximately 250,000-300,000 atoms, with total system size dependent on the specific receptor-G protein complex dimensions and corresponding membrane requirements.

All MD simulations were performed using AMBER20^65^ under NPT ensemble conditions at 310K and 1 bar. The FF19SB^66^ force field was employed for protein parameters, while lipid21^67^ and GAFF^68^ force fields described lipid and ligand interactions, respectively. The OPC^69^ water model was used for solvation, with long-range electrostatic interactions calculated via particle mesh Ewald. A 9 Å van der Waals cutoff was applied throughout all simulations.

System preparation involved energy minimization followed by 47.5 ns of equilibration. Energy minimization was performed using 2,500 steepest descent steps followed by 2,500 conjugate gradient steps. Temperature equilibration employed the Langevin thermostat (γ = 1.0 ps⁻¹) with gradual heating to 310 K over two consecutive 2.5 ns NVT phases using 1 fs timesteps. Pressure equilibration utilized the Berendsen barostat (τ_p_ = 1.0 ps) through a series of NPT phases: an initial 2.5 ns phase at 1 fs timestep, followed by two consecutive 10 ns phases at 2 fs timestep, and a final 20 ns phase at 2 fs timestep with restraints applied only to protein and ligand atom. Harmonic restraints were systematically reduced across the equilibration stages. Lipid restraints decreased progressively to 2.5, 2.5, 1.0, 0.5, 0.1, and 0.0 kcal/mol/Å² across the six equilibration phases. Protein and ligand restraints followed a similar pattern: 10.0, 5.0, 2.5, 1.0, 0.5, and 0.1 kcal/mol/Å². Production simulations were conducted for 1,000 ns per replica with five independent replicates per system, using 2 fs timesteps and coordinates saved every 1,000 ps.

Structural dynamics analyses were performed on simulation trajectories using AmberTools23 CPPTRAJ^70^ for trajectory processing, imaging, and alignment. The simulation trajectories were aligned to the post-minimization starting structure using Cα atoms of residues (MT1 systems: S28–R54, F65–V84, S103–Y128, L145–N162, I189–Q216, F234–G258 and V278–Y295; and MT2 systems: A42–V65, F78–I101, A115–Y139, P158–P174, Y200–A230, F257– L272 and F290–I306) of the 7-transmembrane bundle as the reference framework. Root-mean-square deviation (RMSD) and root-mean-square fluctuation (RMSF) calculations were conducted relative to these aligned post-minimization structures to assess structural stability and regional flexibility throughout the simulation period.

For interaction energy analysis, noncovalent interaction energies between individual receptor residues and the G protein α5 helix were calculated using the NAMDenergy utility^71,72^ (v1.6) distributed with VMD 1.9.3 ^73^. The energy calculations employed a 9 Å cutoff distance with a 7.5 Å van der Waals smoothing switch distance. Energy analyses were performed on every 10th frame of the trajectories (100 frames total from each 1000-frame simulation), corresponding to a 10 ns sampling frequency, across 3 independent replicas of 1 μs MD simulations. Results are reported as mean ± standard deviation values calculated across all replicas to ensure statistical robustness of the findings.

### Statistical analysis

All data are presented as the mean ± standard error of the mean (SEM). Statistical significance was assessed with paired t-test, and two-tailed p-values are reported (Fig. 1c–f, 4d, 5c), or multiple t-tests with Holm–Šidák correction for multiple comparisons (Fig. 2h) and exact p-values are reported throughout. One-way ANOVA followed by Bonferroni’s multiple comparison test (Fig. 5b, 6c) or two-way ANOVA followed by the Holm–Šidák test (Fig. 2d) was used as appropriate. For unpaired comparisons in ex vivo and in vivo experiments, Welch’s correction was applied, and one-tailed p-values are reported (Figures 1h, j, m). P-values < 0.05 were considered statistically significant, and exact p-values are indicated in the figures. All analyses were performed using GraphPad Prism versions 6 and 8 (GraphPad Software Inc., San Diego, CA). Multiple t-tests with Holm–Šidák correction for multiple comparisons and exact p-values are reported throughout. A p-value < 0.05 was considered statistically significant.

## Acknowledgments

We are grateful to Dr. Nevin Lambert (Augusta, Georgia, USA) for sharing some reagents for the G protein recruitment assay, Akihide Yoshimi (National Cancer Center, Tokyo, Japan) for sharing some materials of melatonin receptors, and Takashi Yoshimura (Nagoya University, Aichi, Japan) for helpful suggestions. We are grateful to the Institute of Comprehensive Medical Research, Division of Animal Research Promotion Division (Aichi Medical University) for maintaining the mice, providing advanced research promotion for the expression analysis of proteins, and performing measurements of cAMP in *pars tuberalis* samples.

## Declarations

### Author Contribution

A.O., H.H.O., K.I., B.M. J.D., M.N., I.G.T., O.N. and R.J. conceived and designed experiments or contributed to critical discussion. A.O., B.M. performed biochemical in vitro experiments including the design of receptor mutants and chimera. K.I. performed in vivo studies. H.H.O., T.K. and K.K. performed structural studies including receptor expression and purification, cryoEM data collection and image processing and model building.

T.K. and K.K. performed G protein and Nb35 expression and purification. R.M. and I.G.T. performed molecular simulation studies. A.K., E.C. performed initial in vitro cAMP studies. A.O., H.H.O., K.I., R.M., B.M. I.G.T. and R.J. analysed data.

A.O., K.I., J.D., I.G.T. and R.J. obtained the funding. A.O., H.H.O., and R.J. wrote the first draft of the manuscript which was edited by K.I., B.M., A.K., E.C., J.D. I.G.T., O.N. and approved by all authors.

### Competing interests

O.N. is a cofounder and scientific advisor for Curreio. All other authors declare no competing interests.

**Extended Data Fig. 1 Cryo-EM analysis of the MT_1_-miniG_s_ complex**

The workflow of Cryo-EM analysis of MT_1_-miniG_s_ complex using RELION 3.1 and cryoSPARC v3.3.

**Extended Data Fig. 2 Cryo-EM analysis of the MT_2_ chimera-miniG_s_ complex**

The workflow of Cryo-EM analysis of MT_2_ chimera-miniG_s_ complex using RELION 3.1, RELION 4.0 and cryoSPARC v3.3.

**Extended Data Fig. 3 Structural comparison of the MT_1_-miniG_s_ and MT_2_-MT_1_-miniG_s_ complexes**

a-e, Structural comparison of MT_1_-miniG_s_, MT_1_-G_i_ (PDB 7DB6) and crystal structure of MT_1_ (PDB 6ME2), on (a) overall receptor, (b) ligand binding site, (c) PIF motif, (d) N(D)RY motif and (e) Na^+^ binding site and NPxxY motif.

f-j, Structural comparison of MT_2_-MT_1_-miniG_s_, MT_2_-G_i_ (PDB 7VH0) and crystal structure of MT_2_ (PDB 6ME9), on (f) overall receptor, (g) ligand binding site, (h) PIF motif, (i) N(D)RY motif and (j) Na^+^ binding site and NPxxY motif.

k-m, Structural comparison of G_s_ protein among MT_2_-MT_1_-miniG_s_, β_2_AR-G_s_ (PDB 3SN6) and GDP-bound inactive state (PDB 6EG8), on (k) overall region, (l) α5 helix region and (m) GDP/GTP binding site.

**Extended Data Fig. 4 Structural comparison with GPR174-G_s_ complex and 5HT_4_-G_s_/-G_i_ complex**

a, Structural comparison between MT_1_-miniG_s_ and GPR174-G_s_ (PDB 7VX3).

b, Structural comparison among all the structures of GPR174-G_s_ (PDB 7VX3, 8KH5 and 8IZB).

c, Structural comparison between MT_1_-miniG_s_, GPR174-G_s_ (PDB 7VX3), 5HT_4_-G_s_ (PDB 7XT9) and 5HT_4_-G_i_ (PDB 7XTA).

**Extended Data Fig. 5 Structural comparison of GPCR-G_s_/-G_q_ and GPCR- Gi/-Gq complex with the same receptor**

a, Structural comparison between NK1R-G_s_ (PDB 7P02) and NK1R-G_q_ (PDB 7P00, primary G_q_ coupling).

b, Structural comparison among GPCR-G_i_/-G_q_ complexes with the same receptor; ETBR-G_i_ (PDB 8IY5, primary coupling)/-G_q_ (PDB 8HCX), GASR-G_q_ (PDB 7F8W, primary coupling)/-G_i_ (PDB 7F8V), GHSR-G_q_ (PDB 7W2Z, primary coupling)/-G_i_ (PDB 7F9Y), MRGPRX2-G_q_ (PDB 7S8N)/-G_i_ (PDB 7S8M), NMUR2-G_q_ (PDB 7XK8)/-G_i_ (PDB 7W57), NTSR1-G_i_ (PDB 6OS9, primary coupling)/-G_q_ (PDB 8FMZ), OXYR-G_q_ (PDB 7RYC)/-G_i_ (PDB 7QVM), US28-G_q_ (PDB 7RKF)/-G_i_ (PDB 7RKM) and SSTR2-G_i_ (PDB 7Y24, primary coupling)/-G_q_ (PDB 7Y26).

**Extended Data Fig. 6 Molecular dynamics simulations support differential G protein binding modes and ICL3-dependent Gs coupling selectivity in the melatonin receptors**

a, Representative MD simulation snapshots of the melatonin receptor-G protein complexes. Carbon colours indicate electrostatic energies, stick thickness represents van der Waals interactions with α5 helix. Only residues >1 kcal/mol in TM5/TM6/ICL3 are shown. AlphaFold3-modeled ICL3 (pink) highlights non-conserved hydrophobic residues. TM5 and TM6 are highlighted in darker colour. Key contacts: I210^5.61^ with L348^G.H5.20^, F354^GH5.26^, and L353^GH5.25^ (Gi deep binding) versus V217^ICL3^ with L388^G.H5.20^ (G_s_ shallow binding). The MT2-Gs test model shows compromised interactions due to alanine substitutions.

b, Root mean square fluctuation (RMSF) values of the G protein α5 helix across different melatonin receptor-G protein complexes.

c, G protein α5 helix flexibility (RMSF).

d, ICL3-α5 helix interaction energies. Data in b-d: mean ± SEM, 5×1μs simulations.

