## Supplementary material for "Structural basis and physiological significance of non-canonical Gs coupling to the prototypical Gi-coupled melatonin MT_1_ receptor": Supp and Extended figures

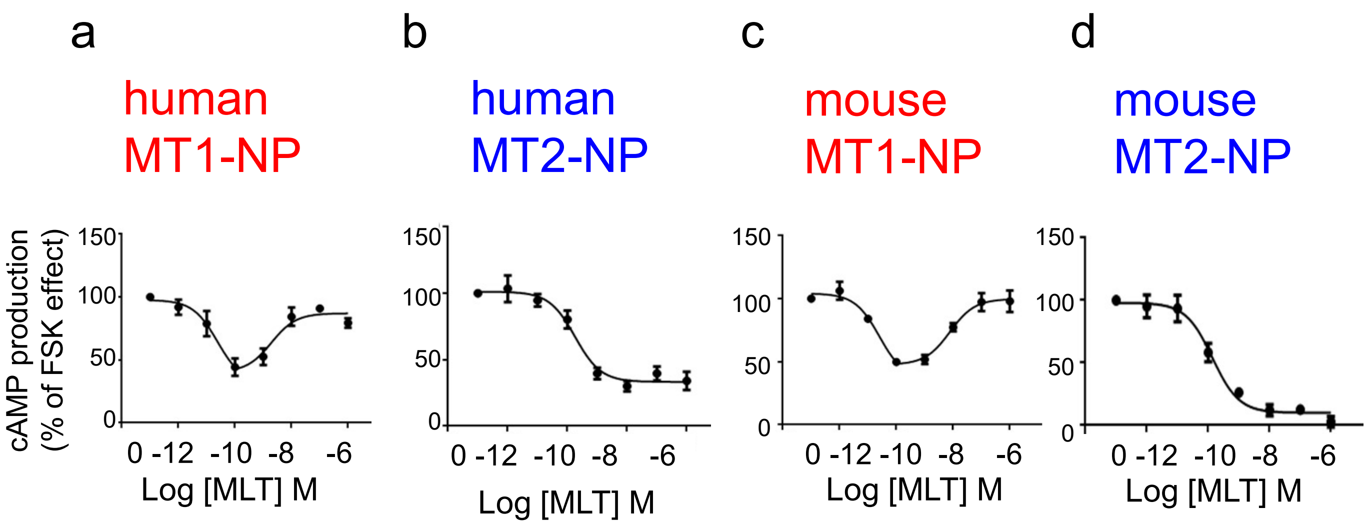


**Supplementary Fig.S1. Dose–response analysis of melatonin-induced cAMP production via C-tail tagged human or mouse melatonin receptor.**

Dose–response analysis of melatonin-induced cAMP production in HEK293T cells expressing Flag-tagged human MT1-NP (a), human MT2-NP (b), mouse MT1-NP (c) and mouse MT2-NP (d) . Cells were treated with forskolin (5 μM) together with the indicated concentrations of melatonin for 10 min. cAMP levels were monitored using the BRET-based CAMYEL biosensor. Data represent means ± SEM from three independent experiments performed in duplicate.

Supplementary Fig. S1

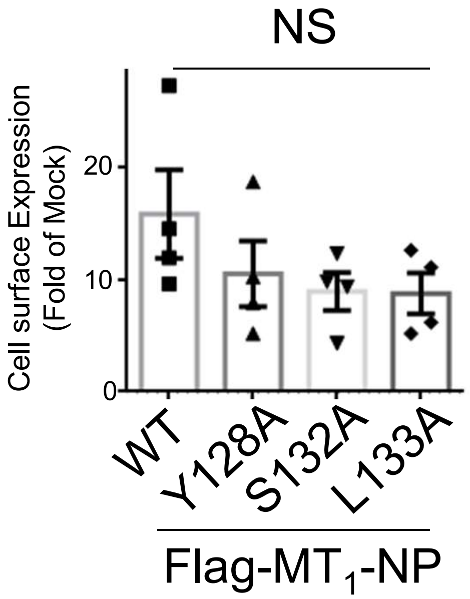


**Supplementary Fig.S2. Cell Surface Expression of MT1 –WT or mutants.**

Cell surface expression of Flag-tagged human WT or mutant MT1 was determined by ELISA. Data represent means ± SEM from three independent experiments performed in duplicate. Statistical analysis was performed by Kruskal-Wallis test followed by Dunn’s multiple comparison test. NS; $p > 0.05$

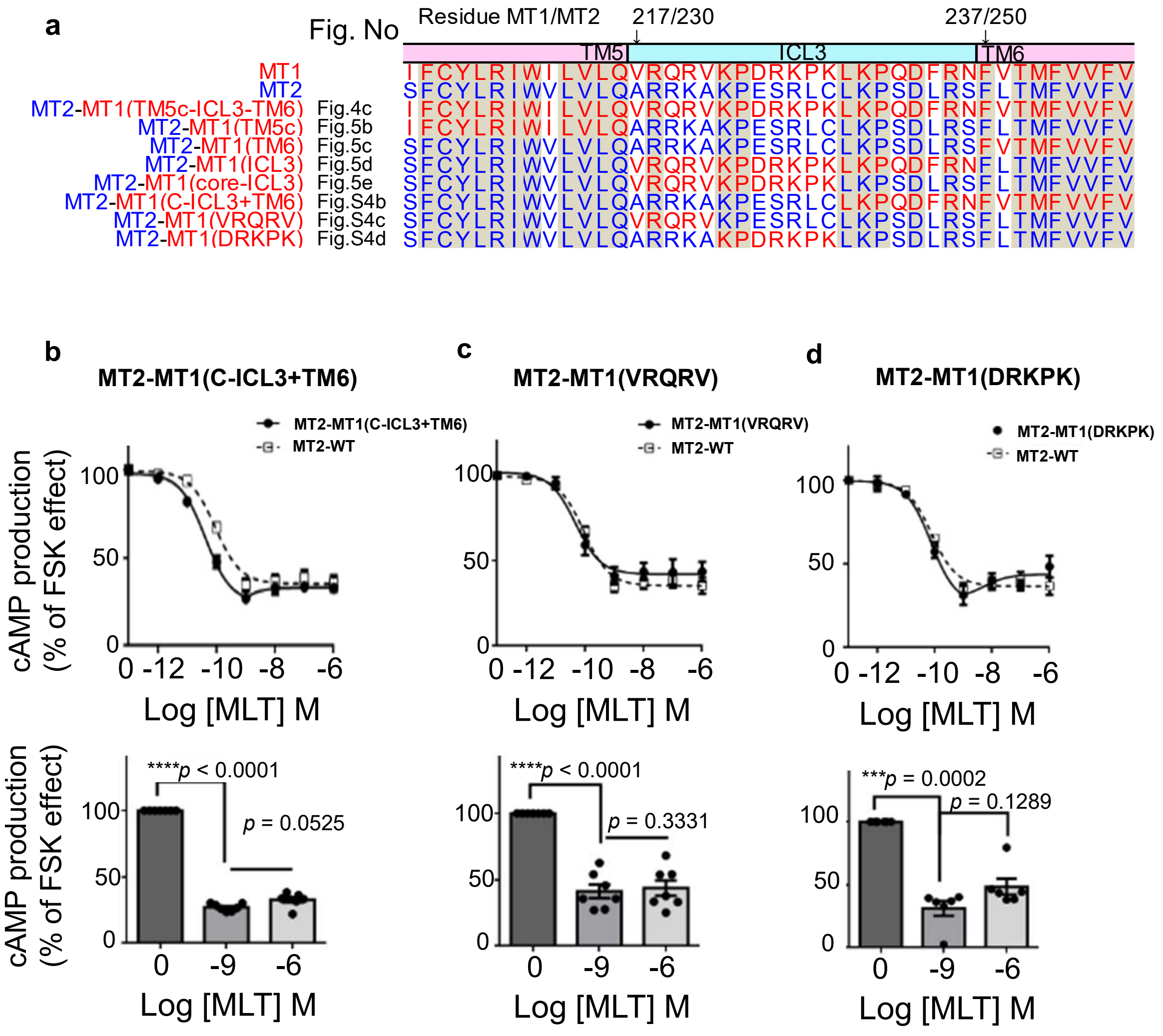


**Supplementary Fig.S3. Melatonin-induced cAMP response of MT2-ICL3 chimera.**

(a) Amino acid sequences alignment of MT2 chimeric receptors (b-d) (upper) Melatonin-induced cAMP production measured by CAMYEL-BRET sensor in transfected HEK293T cells. (lower) Statistical analysis between MLT 0nM and 1nM and 1uM. Data represent means ± SEM from 6 independent experiments performed in duplicate. Statistical analysis was performed by One-Way ANOVA followed by Bonferroni's multiple comparison test.

Supplementary Fig. S3

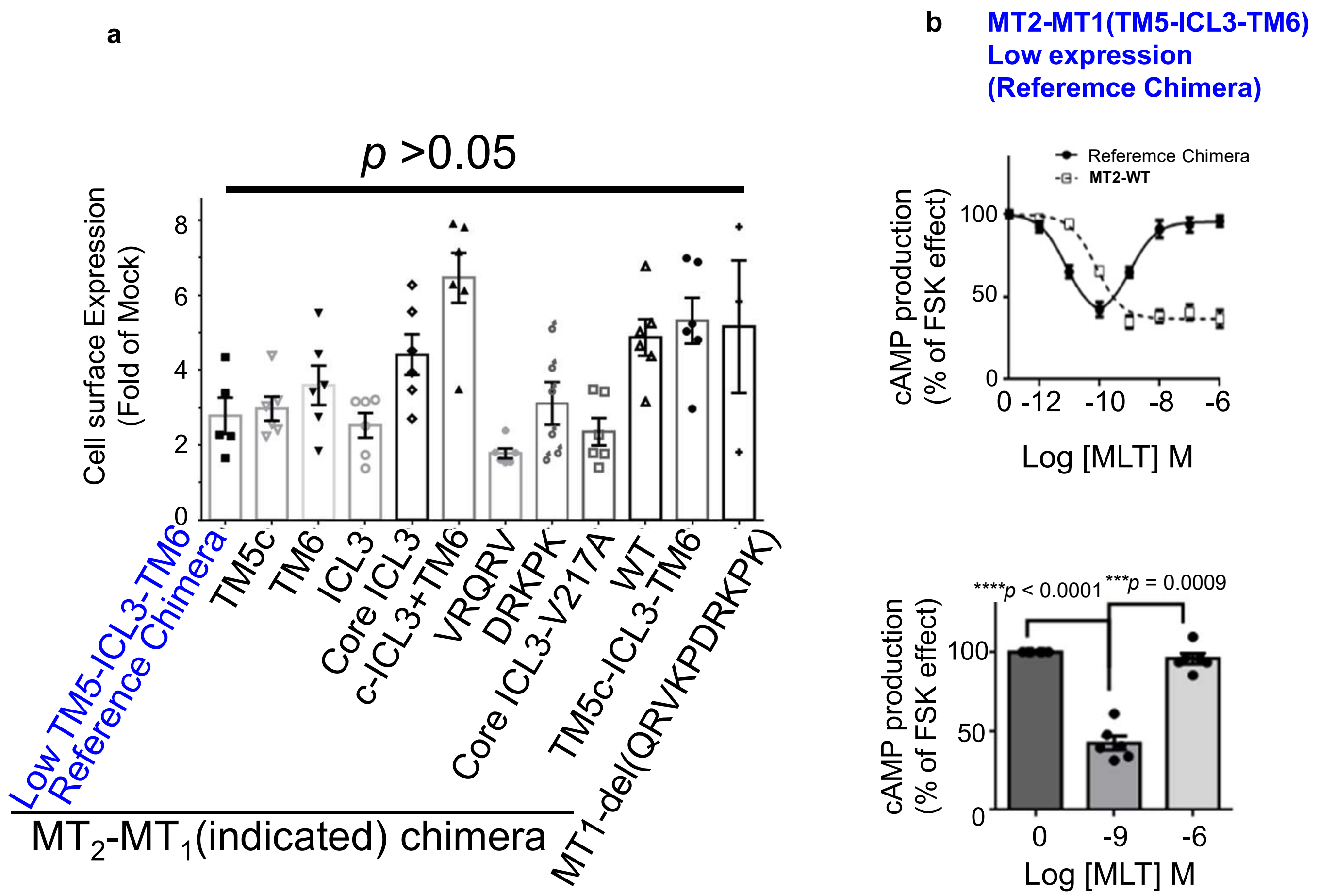


**Supplementary Fig.S4. Cell Surface Expression of MT2 -MT1 Chimera protein and MT1-deletion mutant.** (a) Cell surface expression of Flag-tagged human MT2-WT or chimera MT2 or MT1-deletion mutant were determined by ELISA. The MT2-MT1(TM5-ICL3-TM6) chimera expressed at low levels was used as a reference. Data represent means ± SEM from at least three independent experiments performed in duplicate. Statistical analysis was performed by Kruskal-Wallis test followed by Dunn’s multiple comparison test compared with reference chimera. (b) Concentration-response curve of reference chimera used in panel (a) (MT2-MT1(TM5-ICL3-TM6)).

| System | 7TM Cα RMSD, Å | 7TM Cα RMSF, Å | ICL3 Cα RMSD, Å | ICL3 Cα RMSF, Å | α5 Cα RMSD, Å | α5 Cα RMSF, Å |
|---|---|---|---|---|---|---|
| **MT1_free** | 1.30±0.05 | 0.74±0.03 | 7.54±0.66 | 4.14±0.28[a] | - | - |
| **MT2_free** | 1.20±0.17 | 0.78±0.05 | 5.32±0.65 | 3.84±0.53[b] | - | - |
| **MT1+Gi** | 1.48 ± 0.08 | 0.77 ± 0.05 | 5.98 ± 0.47 | 3.53 ± 0.35[a] | 3.95 ± 0.48 | 2.27 ± 0.27 |
| **MT1+Gs** | 1.36 ± 0.07 | 0.69 ± 0.04 | 7.67 ± 0.59 | 3.28 ± 0.38[a] | 4.17 ± 0.28 | 2.55 ± 0.24 |
| **MT2+Gi** | 1.33 ± 0.13 | 0.76 ± 0.07 | 4.79 ± 0.69 | 2.90 ± 0.38[b] | 3.42 ± 0.41 | 2.27 ± 0.25 |
| **MT2_MT1 (TM5-ICL3-TM6)+Gs** | 1.40 ± 0.08 | 0.73 ± 0.06 | 6.15 ± 0.42 | 3.39 ± 0.35[c] | 5.00 ± 0.43 | 2.76 ± 0.37 |
| **MT2+Gs_testmodel** | 1.40 ± 0.11 | 0.73 ± 0.07 | 7.97 ± 0.86 | 4.68± 0.81[b] | 6.03 ± 0.67 | 3.19 ± 0.35 |

**Supplementary Table 1. Root-mean-square deviation (RMSD) and root-mean-square fluctuation (RMSF) data for all simulated melatonin receptor systems**. Mean ± SEM calculated from three 1 μs replicas for MT1_free and MT2_free, and five 1 μs replicas for MT1+Gi, MT1+Gs, MT2+Gi, MT2_MT1(TM5-ICL3-TM6)+Gs, and MT2+Gs_testmodel. RMSD is measured against the starting point of simulations (after minimization) by Cα atoms of 7-transmembrane bundle residues for alignment. RMSD 7TM-Cα: 7-transmembrane bundle residues - MT1 systems (S28–R54, F65–V84, S103–Y128, L145–N162, I189–Q216, F234–G258, V278–Y295), MT2-based systems (A42–V65, F78–I101, A115–Y139, P158–P174, Y200–A230, F257–L272, F290–I306), RMSD ICL3: intracellular loop 3 residues (V217–P231[a], A230–P244[b], V230–P244[c]), RMSD Alpha5: G protein α5 helix residues (T327–F354 for Gi systems, D368–L394 for Gs systems). RMSF for melatonin receptor regions is measured upon alignment to the average frame of simulations by Cα atoms of 7-transmembrane bundle residues, with regional measurements conducted using the same structural definitions as RMSD calculations.

Supplementary Table. S1

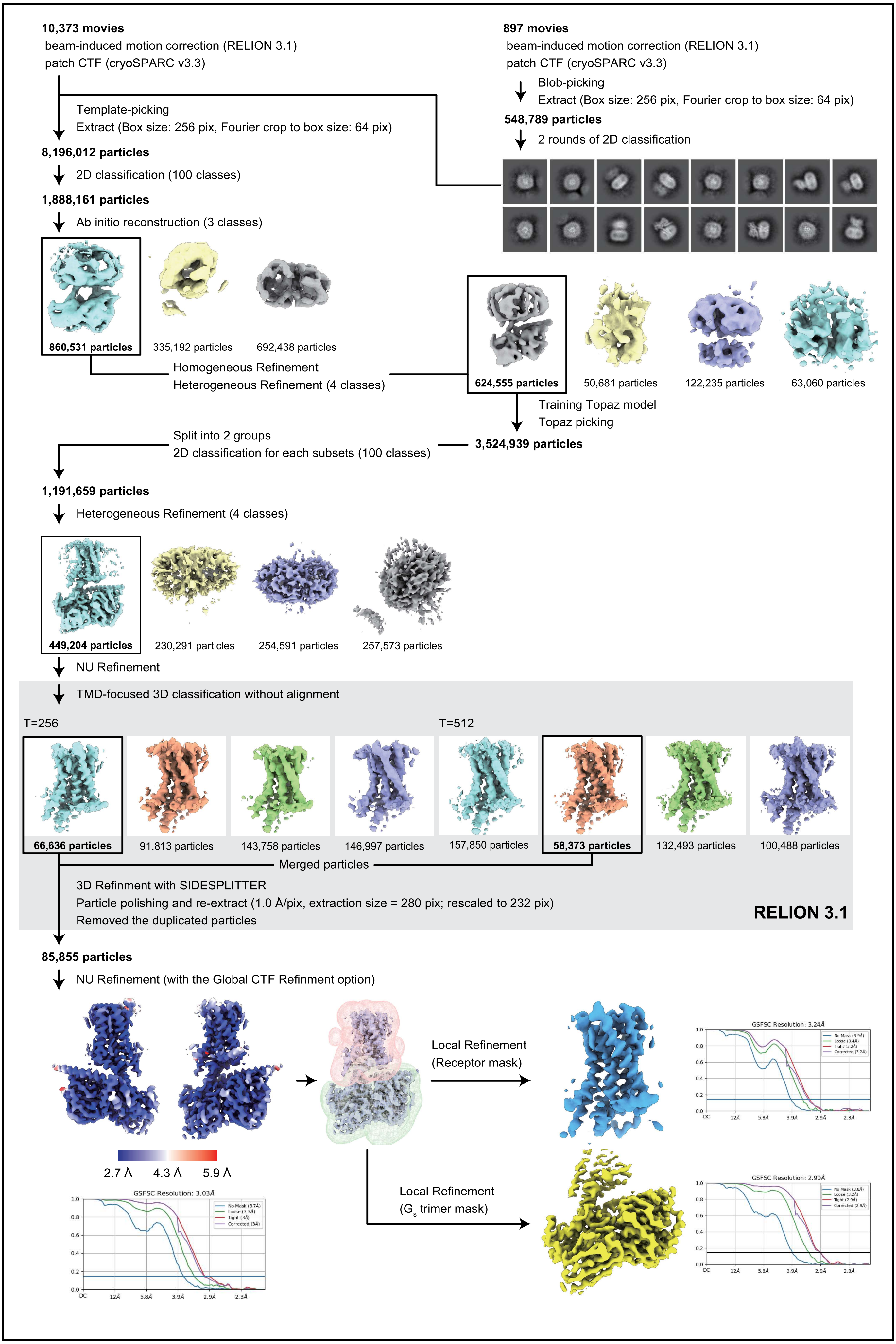


### Extended Data Fig. 1 | Cryo-EM analysis of the $MT_1$-miniG$_s$

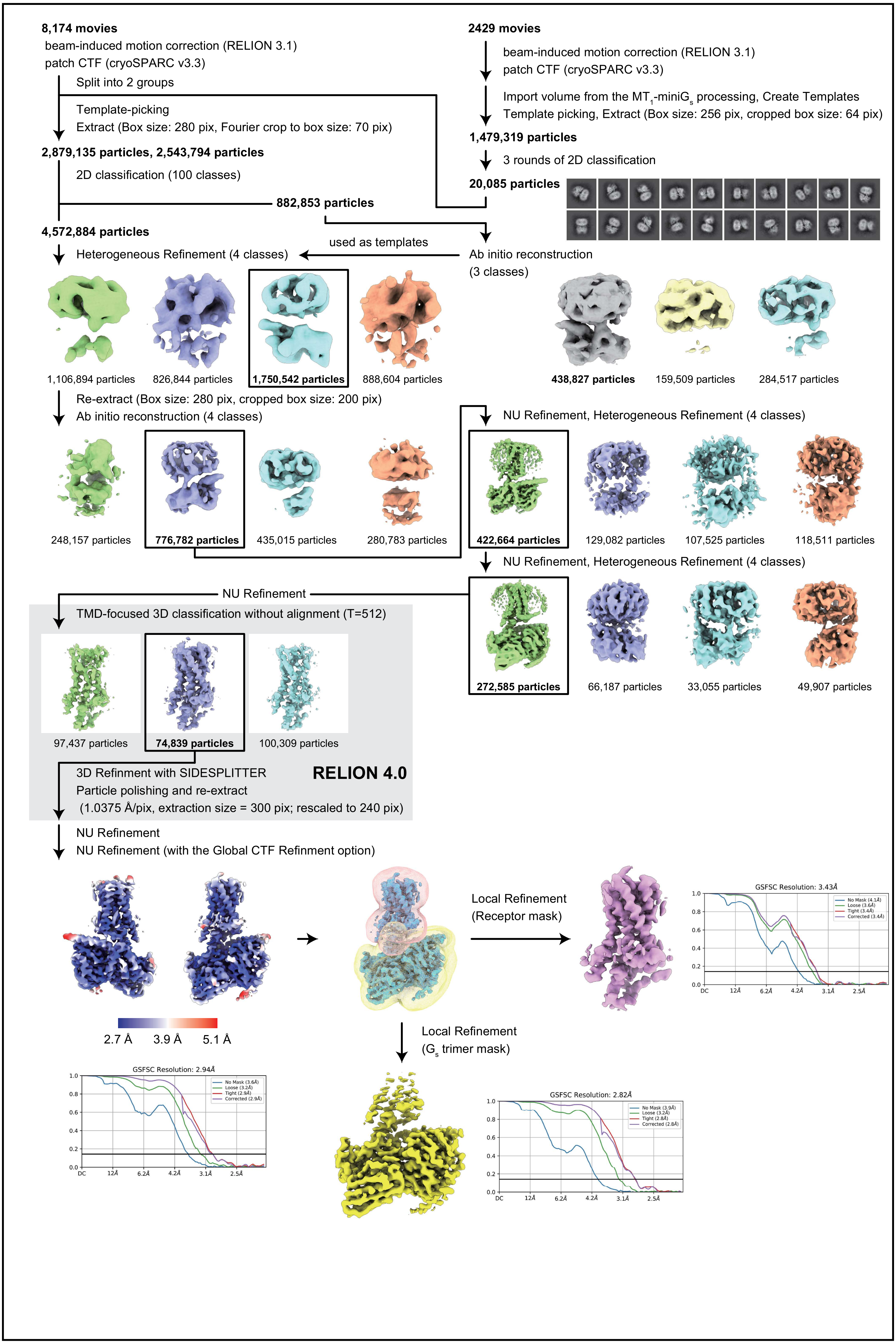


### Extended Figure 2 | Cryo-EM analysis of the $MT_2$ chimera-$miniG_s$

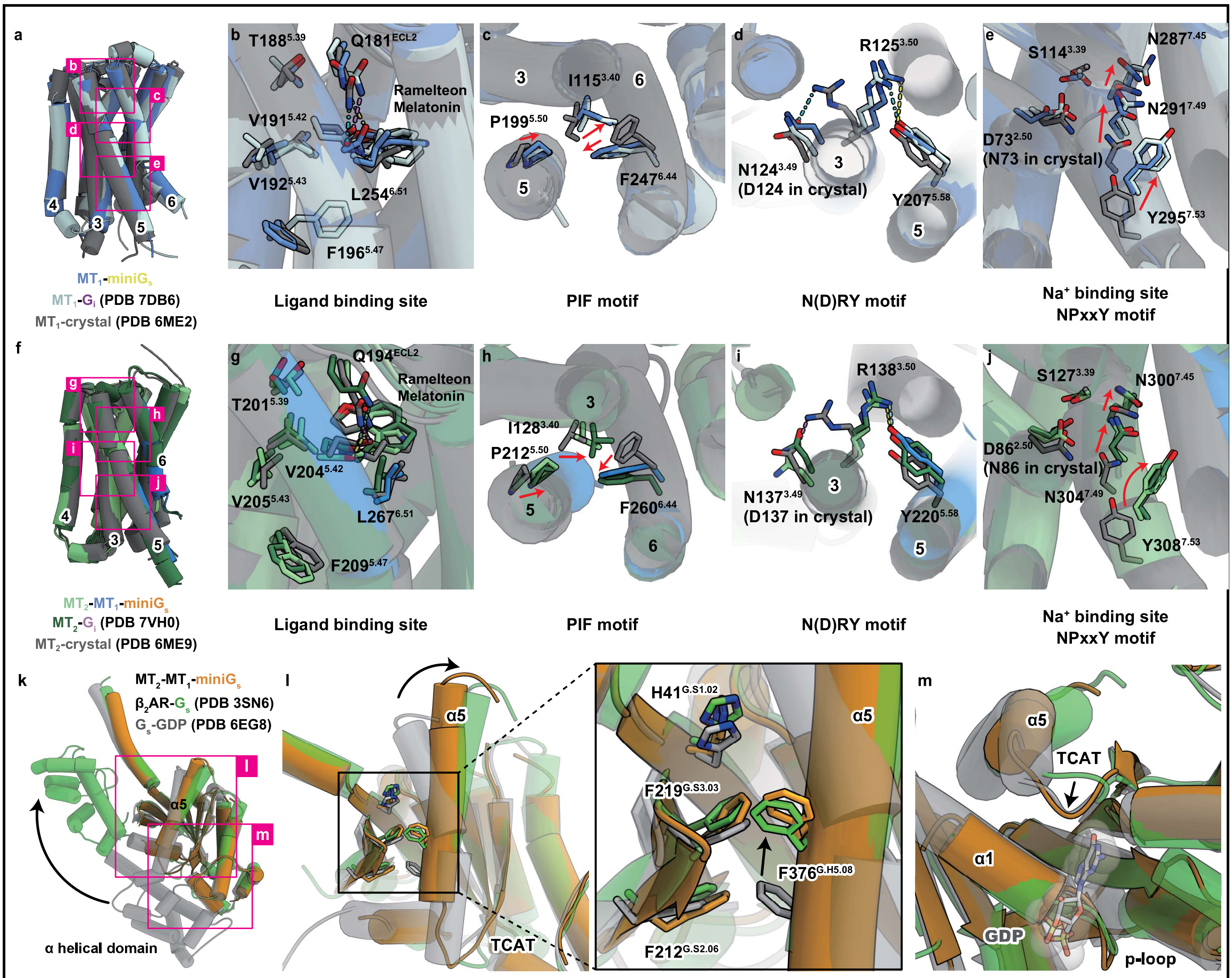


**Extended Figure 3 | Structural comparison among $MT_{1/2}$-$G_{s/i}$ complexes**

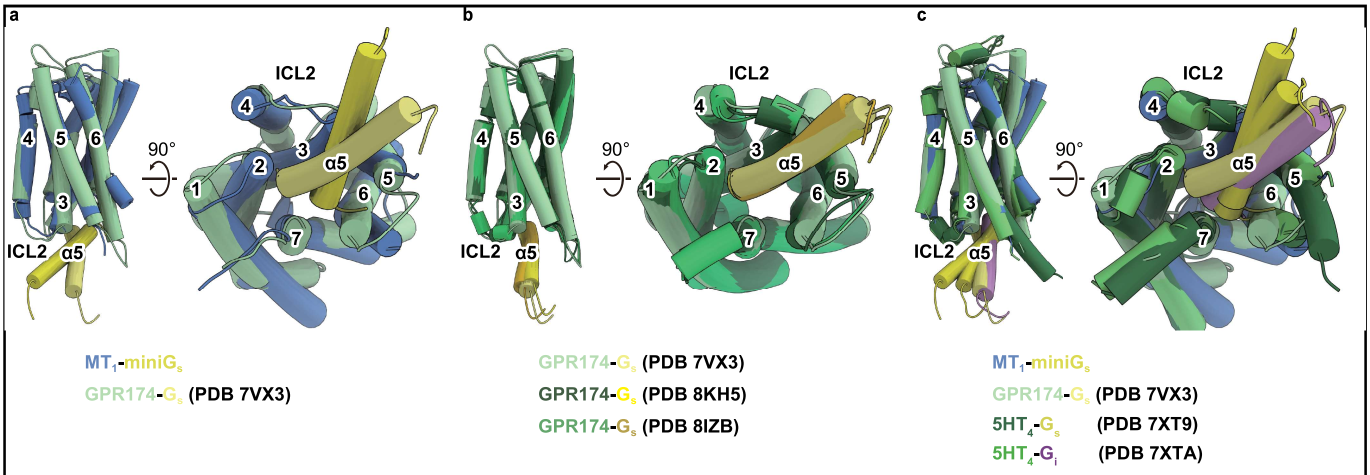


**Extended Figure 4 | Structural comparison with GPR174-G$_s$ complexes**

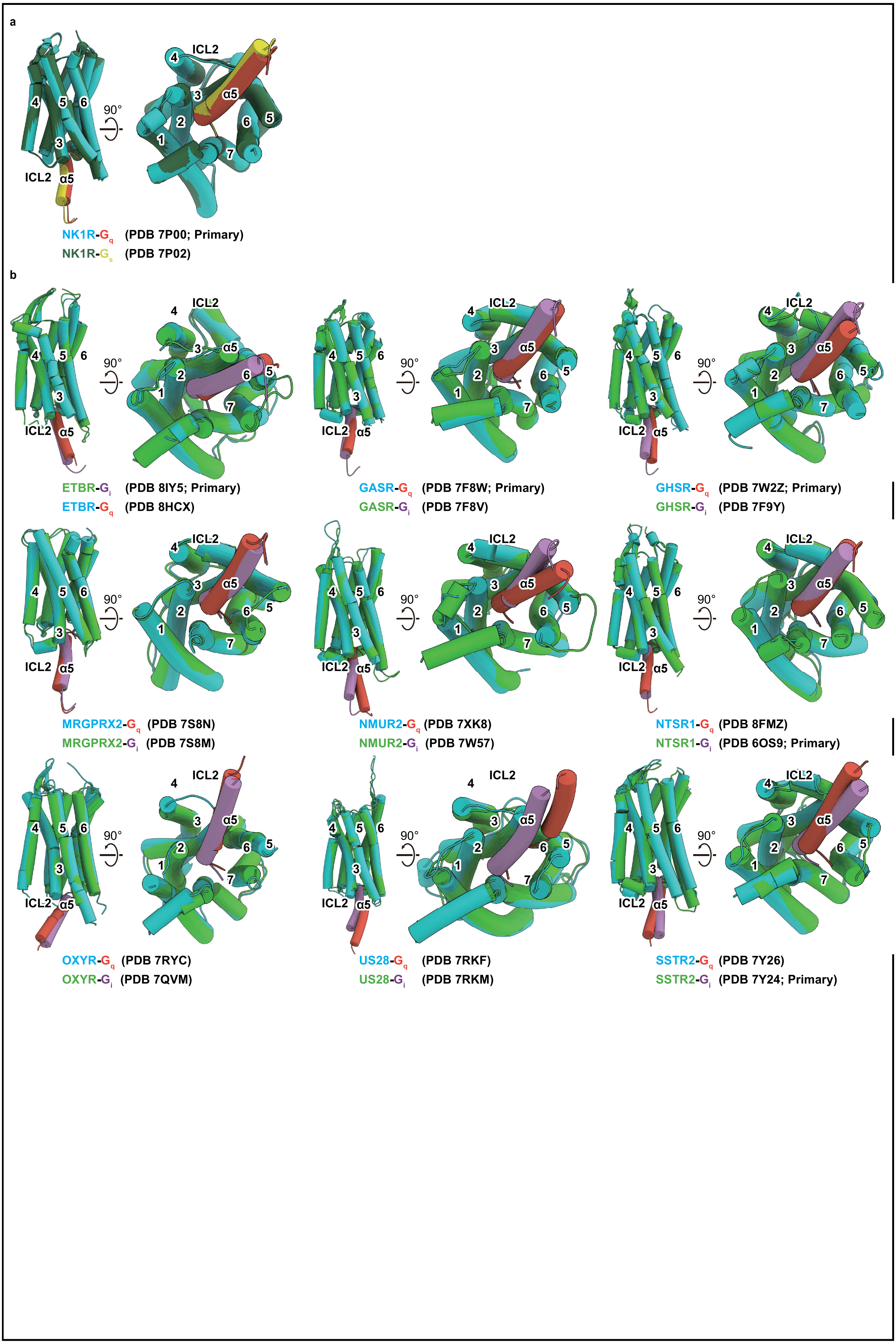


**Extended Figure 5 | Comparison with $G_q$-$G_s$ complexes and $G_q$-$G_i$ complexes**

a, Gs-Gq complex (NK1R). b, Gi-Gq complex (ETBR, GASR, GHSR, MRGX2, NMUR2, NTS1, OXYR, SSTR2, US28)

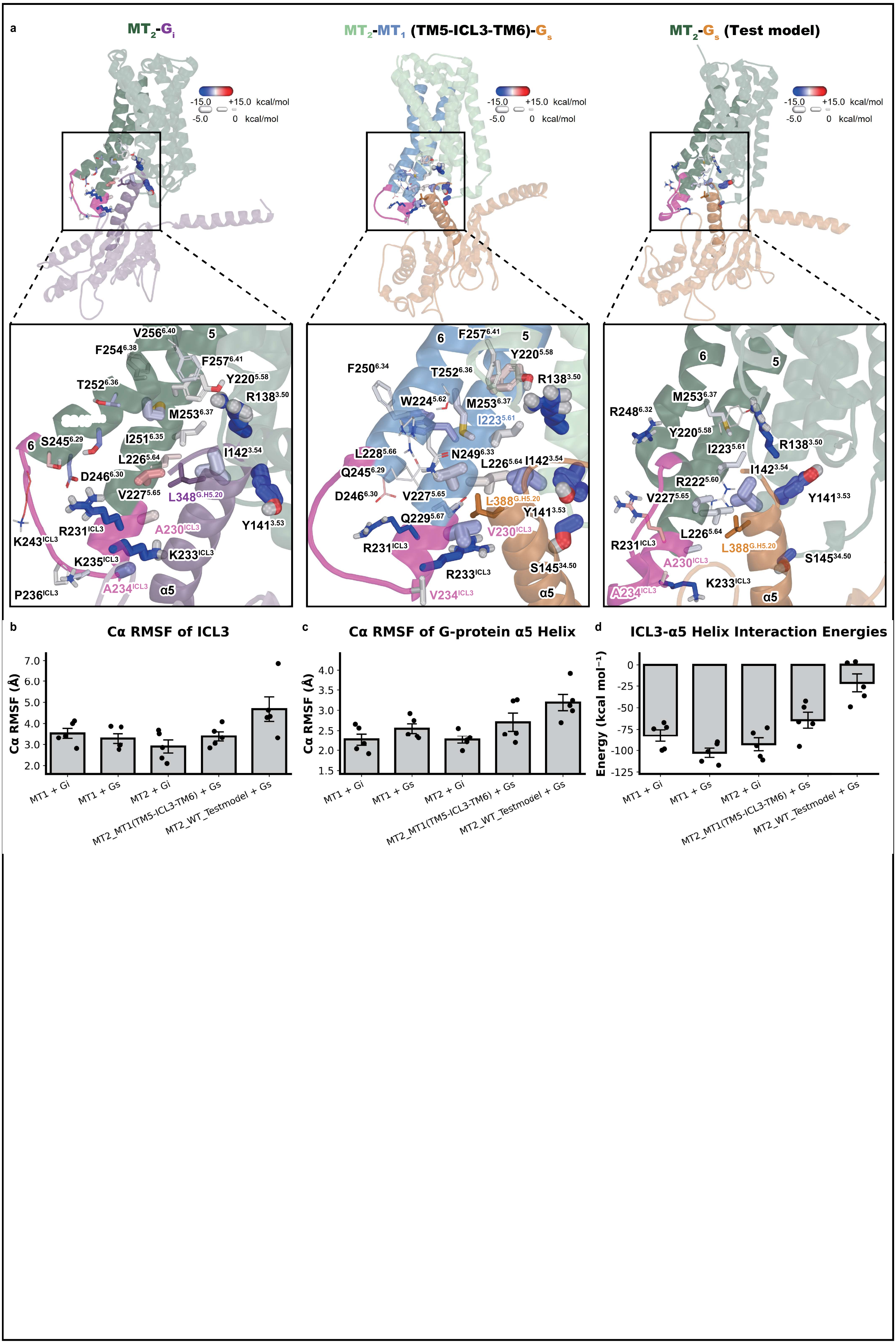


**Extended Figure 6 | Molecular dynamics simulations support differential G protein binding modes and ICL3-dependent**
